# Galectin-3 drives neuroinflammatory amyloid remodeling by stabilizing intermediate Aβ species and altering lysosomal processing

**DOI:** 10.1101/2025.03.17.643790

**Authors:** Lluís Camprubí-Ferrer, Jesús Soldán-Hidalgo, Emil Axell, Marzia Dell’Eva, Marta García-Cruzado, Juan García-Revilla, Yiyi Yang, Rosalía Fernández-Calle, Logan Larlham, Bhanu Chouhan, Rocío Talaverón, Agnes Paulus, Lukas Danielson, Angela Gomez-Arboledas, Oxana Klementieva, Antonia Gutierrez, Javier Vitorica, Malin Wennström, Gunnar K. Gouras, Javier Frontiñán-Rubio, Antonio Boza-Serrano, Gabriel A. Rabinovich, José Luis Venero, Tomas Deierborg

**Affiliations:** Experimental Neuroinflammation Laboratory, Department of Experimental Medical Sciences, Lund University, BMC B11, Lund 221 84, Sweden; Instituto de Biomedicina de Sevilla, IBiS/Hospital Universitario Virgen del Rocío/CSIC/Universidad de Sevilla, Seville, Spain; Departamento de Bioquímica y Biología Molecular, Facultad de Farmacia, Universidad de Sevilla, Seville, Spain; Biochemistry and Structural Biology, Department of Chemistry, Lund University, Lund 223 62, Sweden; Protein Evolution, Department of Experimental Medical Sciences, Lund University, Lund 221 84, Sweden; Medical Microspectroscopy Lab, Department of Experimental Medical Science, SRA: NanoLund, Multipark, Lund University, BMC B10, 221 84, Lund, Sweden; Department of Biología Celular, Genética y Fisiología, Facultad de Ciencias, Instituto de Investigación Biomédica de Málaga-IBIMA Plataforma Bionand, Universidad de Málaga, Malaga, Spain; Centro de Investigación Biomédica en Red sobre Enfermedades Neurodegenerativas (CIBERNED), Madrid, Spain; Cognitive Disorder Research Unit, Department of Clinical Sciences Malmö, Lund University, Malmö, 214-28, Sweden; Department of Experimental Medical Science, Experimental Dementia Research Unit, Lund University, Lund, Sweden; Oxidative Stress and Neurodegeneration Group, Faculty of Medicine Universidad de Castilla-La Mancha, Ciudad Real 13071, Spain; Glycoimmunology Laboratory, CaixaResearch Institute, 08022 Barcelona, Spain; Glycomedicine Laboratory, Institute of Biology and Experimental Medicine, C1428 Ciudad de Buenos Aires, Argentina; Faculty of Exact and Natural Sciences, University of Buenos Aires, C1428 Ciudad de Buenos Aires, Argentina

**Keywords:** Alzheimer’s disease, neuroinflammation, microglia, galectin-3, amyloid-β

## Abstract

Neuroinflammation is a hallmark of Alzheimer’s disease (AD), yet the molecular mediators driving microglial dysfunction and neurotoxicity remain poorly understood. Here, we identify Galectin-3 (Gal3) as a central regulator of plaque-associated microglial responses, linking amyloid-beta (Aβ) aggregation, lysosomal function and plaque-associated neuritic damage. Using postmortem brain tissue from AD patients, we demonstrated that Gal3⁺ microglia are selectively enriched around highly immunogenic dense-core plaques, and correlate with increased LAMP1^+^ dystrophic neurites. Notably, Gal3 was also detected in extra-microglial dystrophic structures and in close association with extracellular amyloid fibrils, suggesting a role in both intracellular and extracellular Aβ dynamics. In APP mice, Gal3 deficiency resulted in more compact plaques, reduced neuronal dystrophies and increased TREM2 expression around amyloid plaques, suggesting altered plaque-associated microglial responses. Mechanistically, *in vitro* studies revealed that Gal3 modulates Aβ uptake and its intracellular processing while lysosomal stress conditions showed increased Aβ fibrillation from monomeric species in Gal3-deficient microglia. In parallel, cell-free assays demonstrated that Gal3 directly interacts with Aβ and selectively inhibits secondary nucleation, thereby stabilizing intermediate assemblies associated with increased neurotoxic potential. Finally, *in silico* and transcriptomic analysis revealed that Gal3 interacts with key immune receptor patterns, having leucine-rich repeats as well as immunoglobulin-like domains. Moreover, Gal3 favors pro-inflammatory microglial programs, while its deletion suppresses type I interferon and microglial neurodegenerative (MGnD) signatures. Together, these findings position Gal3 as a central regulator of amyloid aggregation, lysosomal dysfunction, and microglial activation, driving a neurotoxic phase of AD and highlighting a potential therapeutic window for intervention.

**Significance Statement:** Neuroinflammation and amyloid-beta (Aβ) plaques drive Alzheimer’s disease progression, but the molecular bridges between plaque formation, microglial dysfunction, and neurodegeneration remain unclear. This study identifies Galectin-3 (Gal3) as a pivotal regulator at this interface. We demonstrate that Gal3 is enriched around amyloid dense-core plaques in human brains and directly shapes Aβ aggregation into highly neurotoxic intermediate structures. Genetic deletion of Gal3 in Alzheimer’s mouse models reduces nerve cell damage and change microglial cells toward a protective, less inflammatory state. By linking extracellular protein aggregation with intracellular lysosomal stress and inflammatory gene expression, these findings establish Gal3 as a major driver of Alzheimer’s neurotoxicity and highlight it as a promising therapeutic target

## Introduction

Microglia, the resident immune cells of the central nervous system (CNS), are critical for brain development and homeostasis, and play central roles in the pathogenesis of neurodegenerative diseases (1). They mediate neuroinflammation in response to diverse stimuli through a wide repertoire of receptors and signaling pathways (2). One such stimulus is amyloid-beta (Aβ), a highly aggregation-prone peptide that accumulates in the form of plaques in Alzheimer’s disease (AD). Microglia continuously engage with and process diverse Aβ species throughout the course of disease progression (3), and in doing so, adopt distinct transcriptional, epigenetic and transcriptional profiles. In both mouse models and AD patients, microglia have been demonstrated to acquire disease-associated molecular signatures, such as disease-associated microglia (DAM) and microglial neurodegenerative phenotype (MGnD) linked to Aβ pathology (4–6). These activated states exhibit diverse roles in AD, ranging from facilitating Aβ clearance to driving chronic neuroinflammation, underscoring the functional heterogeneity within the microglial population (7). They have been linked to type I interferon (IFN) and tumor necrosis factor-α (TNF-α), which contribute to disease progression (8, 9). Notably, these pathways present critical opportunities and pharmacological windows for therapeutic modulation (10).

Galectin-3 (Gal3; encoded by *LGALS3* gene), the only chimera-type member of the galectin family binds β-galactoside (Galβ1-4-GlcNAc) residues via its carbohydrate recognition domain (CRD) (11). While primarily monomeric, Gal3 oligomerizes upon ligand binding, enabling crosslinking functions and contributing to processes such as cell adhesion, migration, inflammation, fibrosis, and host responses to infection, cancer, and tissue injury (12–14). Although Gal3 is expressed in several cell types, it becomes highly enriched in microglia within damaged brain regions, where Gal3 can be released upon microglial activation (15–17). Gal3-dependent microglial activation develops a pro-inflammatory profile, which might contribute to a neurotoxic environment leading to neuronal dysfunction in AD (18, 19). Moreover, intracellularly, Gal3 associates with membranes and organelles, especially lysosomes, where it plays a role in sensing and responding to endolysosomal damage (14, 20, 21).

Our group and others have shown that Gal3 is upregulated in microglia surrounding Aβ plaques and that its deletion reduces plaque burden and inflammation in mouse models (4, 16, 22). We recently identified Gal3 as a potential cerebrospinal fluid biomarker for AD (18) and tau-dependent frontotemporal dementia (23). Importantly, the Gal3-targeting monoclonal antibody TB006 (TrueBinding, Inc.) is under clinical evaluation, showing promising early-phase results in AD patients, further supporting the translational relevance of this pathway (Phase 1b/2a, NCT05074498). It is worth mentioning that a recent proteogenomic analysis of CSF and genetic data from more than 1000 patients suffering late-onset AD and controls identified critical AD-associated immune related proteins including TREM2, APOE, PILRA, CD33, SIRPA and Gal3 (24)). Furthermore, a proteogenomic study in Parkinson’s disease identified Gal3 as having a causative role (alongside APOE) as a converging node of innate immune dysregulation (25), reinforcing its relevance across protein aggregation diseases. More broadly, Gal3 has recently emerged as an essential component of universal transcriptomic signatures of mammalian ageing consistently associated with mortality and multimorbidity (26).

Recent findings demonstrated that microglia actively shape Aβ plaques by seeding, compacting, and remodeling them, driving transitions between distinct morphological states. These actions are mediated by microglial receptors, nucleator factors, and lysosomal degradation capacity (27, 28). However, whether Gal3 contributes to these processes by modulating Aβ aggregation dynamics, lysosomal function, or both remains unclear.

In previous work, we have demonstrated that Gal3 is involved in a detrimental form of microglial activation in AD (16, 18) and binds β-sheet-rich fibrils and alters α-synuclein (αSyn) aggregation and toxicity in PD (29). Moreover, we have identified Gal3 as a TLR4 ligand in microglia, mediating proinflammatory responses (15). Furthermore, we have shown that Gal3 also interacts with Triggering Receptor Expressed on Myeloid cells 2 (TREM2), a glycosylated receptor that drives the DAM program, in plaque-associated microglia (6, 30, 31). Interestingly, *Lgals3* deletion significantly alters TREM2-dependent signaling (16). Altogether, these findings place Gal3 as a central regulator orchestrating microglial sensing, signaling, and structural remodeling in response to Aβ pathology. By linking amyloid conformation with receptor activation, lysosomal integrity, and neurotoxicity, Gal3 emerges as a key integrator of cellular and molecular processes that drive AD progression. Here we hypothesize that Gal3 modulates not only the microglial signaling landscape but also plaque morphology, Aβ aggregation dynamics, and the generation of neurotoxic environments marked by dystrophic neurites. Elucidating this complex molecular interplay between microglial Gal3, Aβ kinetics, and lysosomal function will be critical for defining the mechanisms of neuronal injury and for the subsequent development of microglia-targeted therapies in AD as well as other protein aggregation and neurodegenerative diseases.

## Results

### Galectin-3 predominantly associates with neurotoxic cored plaques in the human brain

Our previous work has shown that Gal3 expression occurs around Aβ plaques, although a detailed characterization of its association with specific plaque types was lacking (16, 18). Here, we sought to determine which amyloid-β (Aβ) plaque subtypes in the human brain are most closely associated with microglial Gal3.

We analyzed postmortem hippocampal tissue from nine human AD subjects (Supplementary Table 1). Gal3-positive (Gal3⁺) microglia were identified by colocalization with the microglial marker Iba1. The presence of Gal3⁺ microglia on Aβ plaques was heterogeneous; we therefore classified plaques as Gal3⁻ or Gal3⁺ (Fig. 1A). We also categorized Aβ plaques into four types based on Methoxy-X04 staining (dye staining fibrillar β-sheet-rich amyloid structures) and Aβ immunoreactivity using MOAB2 antibody (which targets N-terminal Aβ epitopes present in Aβ40 and particularly Aβ42 species), without formic acid epitope exposure. The four defined types were: inert (Methoxy⁺/MOAB2⁻), compact (Methoxy⁺/MOAB2⁺ with complete colocalization), filamentous (Methoxy⁺/MOAB2⁺ with diffuse amyloid filaments), and cored (dense Methoxy⁺ core with an MOAB2⁺ halo) (32, 33) (Supplementary Fig. 1A). Notably, diffuse plaques (Methoxy-/MOAB2⁺ with diffuse MOAB2 staining) were very rare across subjects and therefore not quantifiable (Supplementary Fig. 1A). In general, inert plaques were the most prevalent type of Aβ accumulation (38-62%), followed by compact (26-38%), filamentous (8-20%), and cored plaques (2-8%) (Fig. 1B), with some variability across specific brain region.

**Figure 1.**
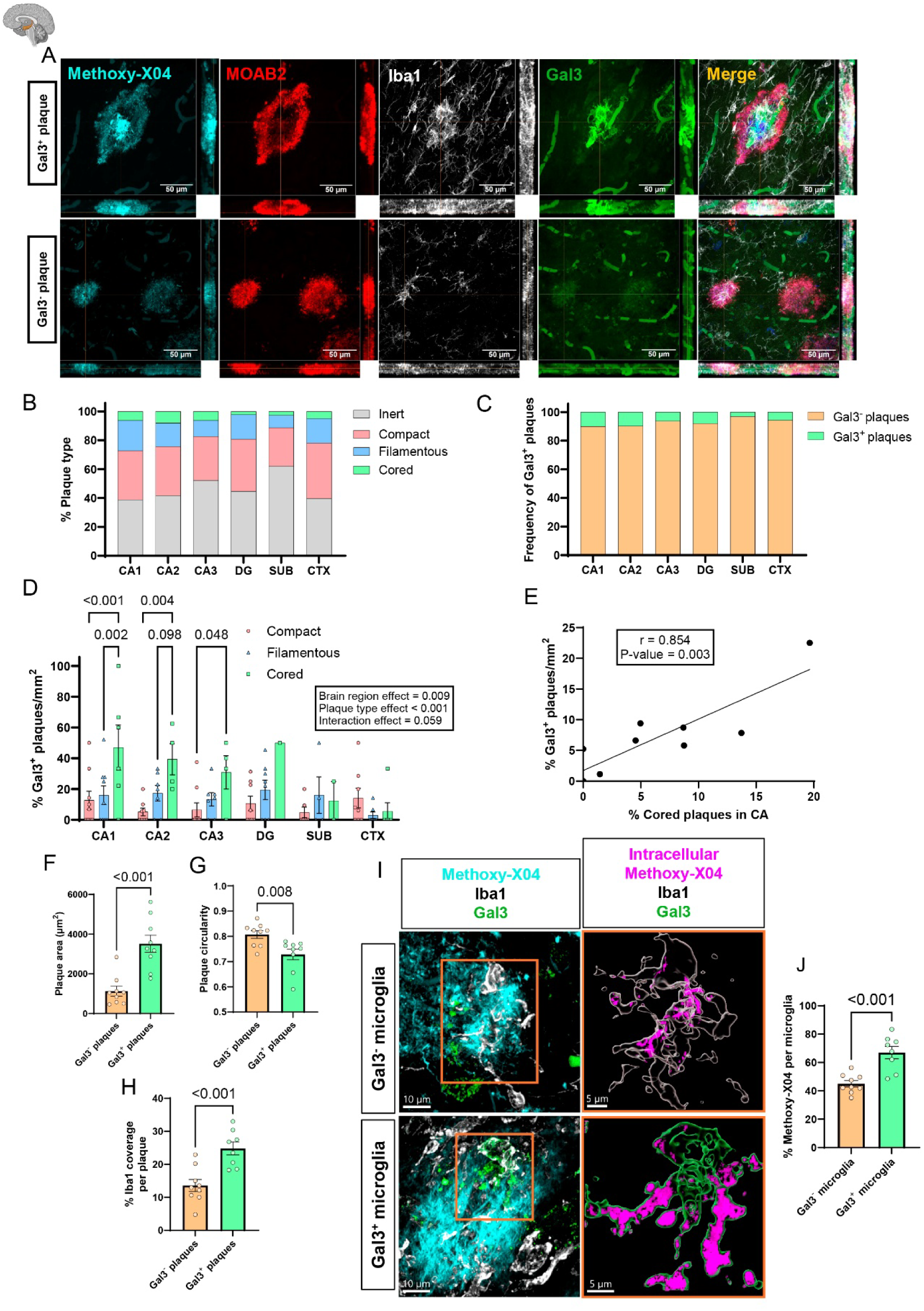
Microglial Galectin-3 selectively associates with cored amyloid plaques and marks phagocytically active states in human AD hippocampus. **A** Representative immunostaining of amyloid plaques showing Gal3^+^ microglia (Gal3^+^ plaque) and Gal3^-^ microglia (Gal3^-^ plaque), stained with amyloid plaque markers Methoxy-X04, MOAB2 (Aβ) and microglial markers Iba1 and Gal3. **B** Average proportions of quantifiable types of plaques across regions. **C** Average proportions of Gal3^-^ and Gal3^+^ plaques across regions. **D** Percentage of Gal3⁺ plaques per patient by region and plaque type. For each plaque type in each region, the percentage of Gal3⁺ plaques was calculated relative to the total number of that plaque type. Only plaque types found ≥2 times per region per patient were included to allow valid percentage calculation. **E** Correlation between the percentage of cored plaques and the percentage of Gal3^+^ plaques found per mm^2^. **F** Average plaque area per patient in Gal3^-^ and Gal3^+^ plaques. **G** Average plaque circularity per patient in Gal3^-^ and Gal3^+^ plaques. **H** Average microglial coverage per patient in Gal3^-^ and Gal3^+^ plaques. **I** Representative 3D reconstruction of Gal3^-^ and Gal3^+^ microglia around a plaque with the amount of intracellular Methoxy-X04 content in each cell. **J** Average intramicroglial Methoxy-X04 per patient in Gal3^-^ and Gal3^+^ microglia. In D, F, G, H, J, values are expressed as individual experimental replicates with mean ±SEM. In D, two-way ANOVA with Tukey’s multiple comparisons was performed. In E, Pearson correlation was performed. In F, G, H and J, unpaired t-test was performed. P-values are expressed with 3 decimals. DG = dentate gyrus, SUB = Subiculum, CTX = Cortex.

We next quantified plaques containing Gal3⁺ microglia (referred as “Gal3⁺ plaques”). Gal3 positivity was observed in a proportion of plaques ranging from 3 to 10% of total plaques, varying by region (Fig. 1C). Inter-individual variability was observed across brain regions (Supplementary Fig. 1B). Notably, Gal3^+^microglia preferentially associated with cored plaques, particularly in the cornu Ammonis (CA) region (Fig. 1D). Similarly, the frequency of cored plaques positively correlated with the number of Gal3^+^ plaques in this region (Fig. 1E). The association with cored plaques might indicate a preference for intermediate stage in amyloid plaque maturation (33). We next examined the morphological characterization of Gal3⁺ plaques. These plaques were significantly larger and exhibited a more irregular shape compared to Gal3⁻ plaques (Fig. 1F, G), in agreement with our prior observation (18). In addition, Gal3⁺ plaques showed nearly twofold increased microglial coverage (Fig. 1H), suggesting enhanced microglial engagement. Consistent with this finding, Gal3⁺ microglia contained increased intracellular amyloid material, as indicated by Methoxy staining (Fig. 1I, J), reflecting elevated phagocytic activity and/or processing of Aβ.

Together, these findings demonstrate that Gal3⁺ microglia are selectively associated with cored plaques, a plaque subtype enriched in structurally diverse and potentially neurotoxic Aβ species (34, 35). Moreover, Gal3 expression is linked to increased microglial engagement and altered plaque morphology, supporting a role for this lectin in shaping microglia-Aβ interactions during specific stages of plaque evolution.

### Galectin-3 promotes plaque expansion and neuritic pathology *in vivo*

Our analysis of human tissue suggested that Gal3 is associated with specific plaque morphologies. To determine whether Gal3 contributes to Aβ plaque architecture *in vivo*, we analyzed 18-month-old APP mice, a well-established mouse model for advanced amyloid pathology.

Consistent with previous reports (16, 22), mice lacking Gal3 exhibited a significantly reduction in overall plaque burden in both cortex and hippocampus, as measured by Thioflavin-S (ThS) and Aβ (BAM10 antibody) staining (Fig. 2A, B). To further assess plaque architecture, we first quantified plaque morphology in the hippocampus (Fig. 2C). Gal3 deletion resulted in a marked reduction in plaque area, circularity, and solidity, indicating altered plaque morphology with a more irregular outline (Fig. 2D). We next assessed Aβ packing within plaques. While ThS staining remained unchanged (Supplementary Fig. 2A), Gal3 deletion increased Aβ staining intensity (Fig. 2D) and the ThS/Aβ ratio (Fig. 2E), consistent with enhanced plaque compaction.

**Figure. 2.**
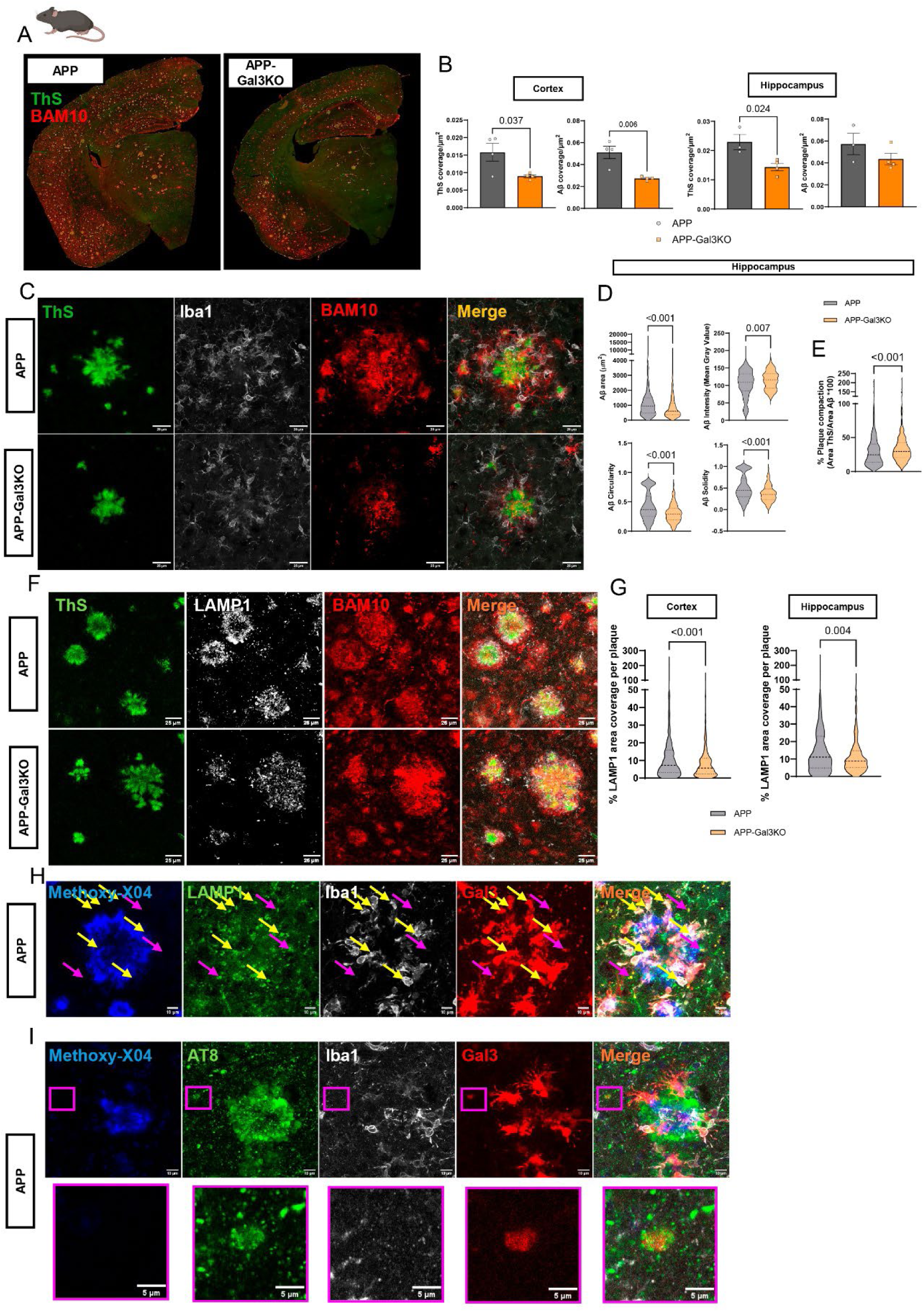
Galectin-3 deletion enhances Aβ plaque compaction and reduces dystrophic neurites. **A** Representative coronal section of APP and APP-Gal3KO mouse brain, stained with amyloid plaque markers Thioflavin-S (ThS), BAM10 (Aβ). **B** Quantification of total plaque burden per mouse (n≥3/group). **C** Representative individual plaque staining of ThS, microglia (Iba1) and Aβ. **D** Morphology parameters of individual plaques by Aβ quantified in hippocampus (n=75 plaques per mouse, ≥4 mice per group). **E** Proportion of plaque compaction measured by Area ThS/Area Aβ*100 in hippocampus (n=75 plaques per mouse, ≥4 mice per group). **F** Representative individual plaque staining of dystrophic neurite marker LAMP1 in APP and APP-Gal3KO mouse brains. **G** Quantification of individual plaque dystrophic neurite content in cortex and hippocampus (n=75 plaques per mouse, 5 mice/group). **H** Representative individual plaque staining of Methoxy-X04, LAMP1, Iba1 and Gal3 in APP mice showing LAMP1^+^Gal3^+^Iba1^-^ dystrophic neurites (pink arrows) as well as LAMP1^+^Gal3^+^Iba1^+^ intra-microglial lysosomal activation (yellow arrows). **I** Representative individual plaque staining of Methoxy-X04, AT8, Iba1 and Gal3 showing AT8^+^Gal3^+^ dystrophic neurites. In B, values are expressed as individual experimental replicates with mean ±SEM. In D, E and G, values are expressed as violin plots. In B, unpaired t-test was performed. In D, E and G, Mann-Whitney test was performed. P-values are expressed with 3 decimals.

These findings suggest that Gal3 promotes the expansion of the Aβ⁺ halo, without substantially affecting the fibrillar core, consistent with the increased plaque size area observed in patients’ Gal3^+^ plaques. Indeed, a vast majority of plaques displayed a ThS/Aβ ratio below 100%, resembling the filamentous and cored plaque structures observed in human samples (Supplementary Fig. 1A). Cortical plaque quantification did not recapitulate these findings (Supplementary Fig. 2B-D), reinforcing the brain region-specific pattern observed in human samples.

Given the established link between plaque structure and toxicity, we next assessed the frequency of lysosomal dystrophies, typically found in dystrophic neurites, which represent a key pathological feature of amyloid plaques. These structures are swollen or aberrant neuronal processes (36) that form around amyloid plaques and contribute to synaptic disruption and neuronal dysfunction (35, 37). These structures can be identified using lysosomal markers such as LAMP1, based on their high accumulation of lysosomal alterations. Interestingly, APP-Gal3KO mice showed a significant reduction of LAMP1 in both cortex and hippocampus (Fig. 2F, G), indicating that Gal3 contributes to neuritic pathology. Notably, Gal3 was detected within a subset of extra-microglial LAMP1 accumulations (Fig. 2H, pink arrows), which were further confirmed to be neuritic dystrophies by colocalization with phosphorylated Tau (AT8) (Fig. 2K). Nevertheless, LAMP1 was also abundantly present in Gal3^+^ microglia (Fig. 2H, yellow arrows), suggesting a relation between Gal3 and lysosomal performance in microglia, however, no changes in phagocytic marker CD68 were observed (Supplementary Fig. 2E,F). Because microglial TREM2 signaling has been implicated in plaque compaction, and neuroprotection (27, 38, 39), we next examined whether Gal3 deletion affected this pathway. APP-Gal3KO mice showed increased TREM2 coverage around plaques (Supplementary Fig. 2E,G). Consistent with this, Gal3KO BV2 microglial cell line displayed increased TREM2 protein levels following Aβ exposure (Supplementary Fig. 2H-J) *in vitro*.

Collectively, these findings indicate that Gal3 deficiency reduces plaque burden, limits Aβ halo expansion, favours plaque compaction and decreases plaque-associated LAMP1 pathology, while being associated with increased TREM2 coverage. Our data supports a hypothesis in which Gal3 contributes to the formation of structurally less compact and more neurotoxic plaque environment, thereby promoting AD progression.

### Galectin-3 localizes to microglial lysosomes and neuritic dystrophic compartments

Given the strong association between cored Aβ plaques and dystrophic neurites (34, 35), we next examined whether Gal3 is localized to these structures in AD patients. LAMP1-Iba1 immunostaining revealed two spatially and cellularly distinct LAMP1⁺ populations: (1) Iba1⁻, extra-microglial LAMP1⁺ structures consistent with dystrophic neurites (Fig. 3A, pink arrows), and (2) Iba1⁺, LAMP1-enriched microglia, corresponding to microglial lysosomal compartments (Fig. 3A, yellow arrows). This distinction is particularly relevant given that Gal3 can be secreted by microglia (15) and subsequently internalized by neurons (29). To confirm the neuronal identity of extra-microglial Gal3⁺ structures, we performed AT8 immunostaining and observed colocalization of Gal3 with AT8⁺ dystrophic neurites outside Iba1⁺ cells (Supplementary Fig. 2K), in line with our mouse model. Quantitative analysis showed that microglial Gal3 levels around plaques positively correlated with the density of LAMP1⁺ dystrophic neurites (Fig. 3B), AT8⁺ neurites (Supplementary Fig. 2L), and plaque core intensity (Fig. 3C). In contrast, Gal3 localized within dystrophic neurites correlated with dystrophy burden but not plaque core intensity (Supplementary Fig. 2M, N), indicating compartment-specific associations with plaque pathology.

**Figure. 3.**
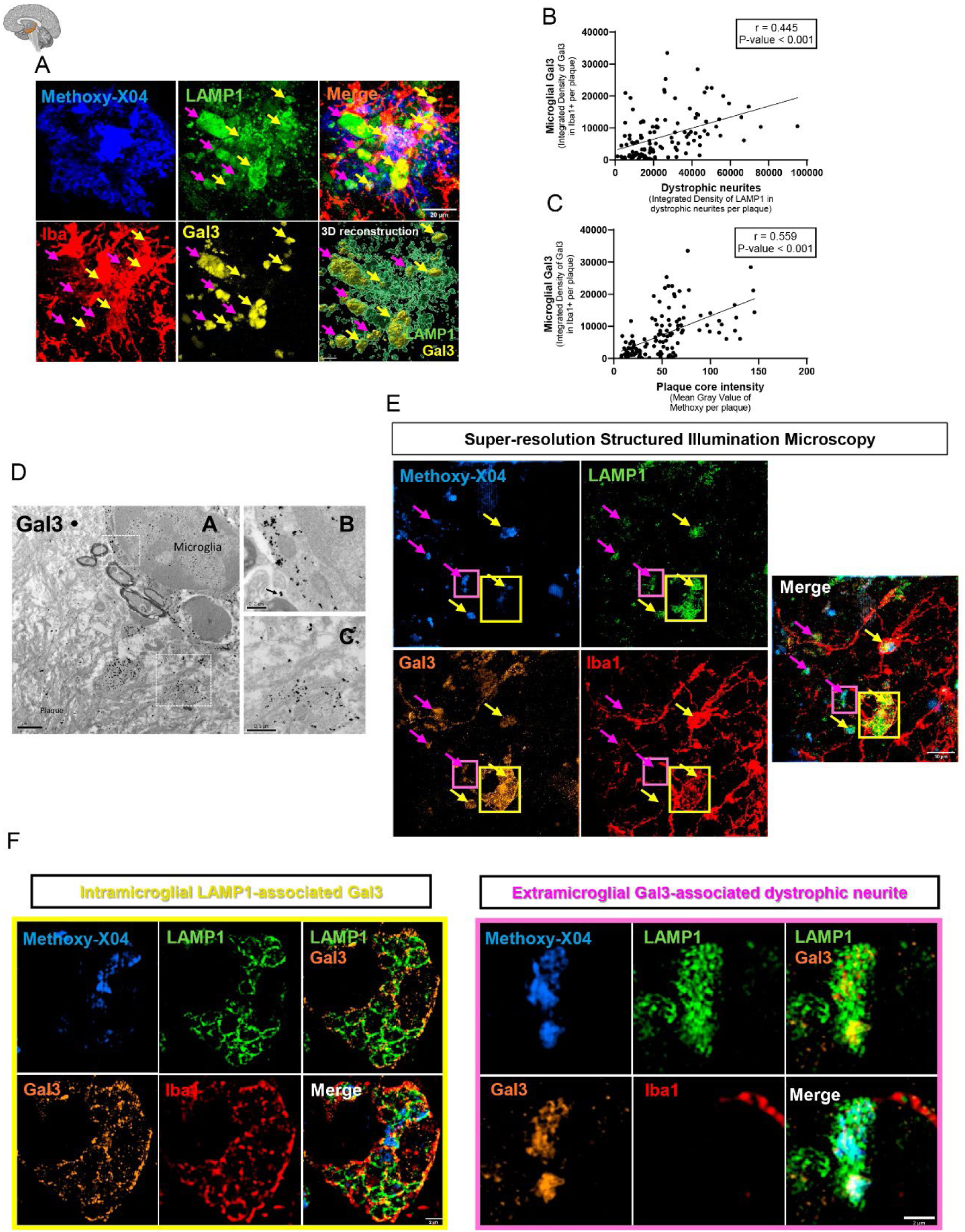
Microglial Galectin-3 colocalizes with lysosome-rich dystrophic neurites and contributes to plaque compaction in Alzheimer’s disease. **A** Representative immunostaining and 3D reconstruction of an amyloid plaque showing dystrophic neurites, stained with Methoxy-X04, LAMP1, Iba1 and Gal3. **B** Individual plaque correlation analysis between dystrophic neurites density and microglial Gal3 density. **C** Individual plaque correlation analysis between plaque intensity and microglial Gal3 density. **D** Gold-immunolabeled electron micrographs of Gal3 in the hippocampus of APP/PS1 mice. A) Immunogold particles specifically labelled microglial cells (Microglia) in the vicinity of amyloid plaques (Plaque). Immunoelectron microscopy confirmed Gal3-labeling also in the extracellular space and in fibrillar aggregates. B) Higher magnification image showing gold particles within the microglial cytoplasm as well as extracellular (arrow). C) Detail image of Gal3 immunolabeling associated with extracellular bundles of plaque-associated amyloid fibrils. **E** Representative immunostaining of N-SIM-reconstruction. **F** Zoomed-in of N-SIM reconstruction showing: Gal3 localized within dystrophic neurites (white arrows), Gal3 associated with non-phagocytic lysosomal compartments (yellow arrows), and Gal3 present in phagocytic lysosomal compartments (orange arrows). In B and C, Pearson correlation was performed.

Next, we sought to determine the origin of neuritic Gal3. We performed immunogold labeling in APP/PS1 mouse brain sections (Fig. 3D). Transmission electron microscopy revealed the presence of Gal3 at the microglial plasma membrane and in the extracellular peri-membranous space, as well as associated with Aβ fibrils within plaques (Fig. 3D), supporting the notion that microglia secrete Gal3 in response to Aβ (16). To further resolve Gal3 localization at the subcellular level, we performed super-resolution SIM imaging. This analysis corroborated the association between intra-microglial Gal3 and lysosomal structures (yellow arrows and square) as well as extra-microglial Gal3 and dystrophic neurites (pink arrows and square) (Fig. 3E,F). Notably, in microglia,

Gal3 appeared as discrete, LAMP1 and/or Iba1 membrane-associated staining surrounding intra-microglial Aβ, while microglia present enlarged lysosomes in a foamy morphology. In extra-microglial compartments such as neurites or peri-plaque space, direct Gal3-Methoxy-X04 can be observed where Gal3 appears as diffused and Methoxy-X04 intensity is attenuated.

Overall, these findings demonstrate that Gal3 localizes to both microglial lysosomal compartments and dystrophic neurites, due to microglial Gal3 release ability, positioning it at the interface between intracellular Aβ processing and neuritic pathology.

### s-μFTIR reveals dynamic changes in Aβ secondary structure during microglial processing

To overcome limitations of fluorescence-based approaches in assessing Aβ aggregation states, we used synchrotron-based Fourier transform infrared micro-spectroscopy (s-μFTIR) to analyze structural changes in Aβ during microglial processing. WT BV2 cells were exposed to fibrillar Aβ (fAβ) for 1, 12 or 24 h, with untreated cells, cell-free fAβ and cells treated with mAβ for 24 h as controls.

Protein secondary structure was assessed by analyzing the amide I region, which reflects β-sheet formations (Supplementary Fig. 3A). Second derivative analysis resolved overlapping peaks in the amide I region, identifying key amyloid structures (Supplementary Fig. 3B) (40–42): β-sheet structures (detected at ∼1629 cm⁻¹), which correspond to intermolecular structures in mature fibrils (43, 44); unordered β-sheets (detected at ∼1639 cm⁻¹) that represent transient early aggregation states (45, 46); 1656 cm^-1^, characteristic of α-helices; antiparallel β-sheets (∼1691 cm⁻¹), which are linked to neurotoxic oligomeric Aβ species (47–50).

As expected, untreated BV2 cells displayed low β-sheet content, whereas, cell-free fAβ showed high levels (Supplementary Fig. 3E). In BV2 cells exposed to fAβ, β-sheet content increased at 1 , consistent with effective internalization, and subsequently decreased by 12 h to levels comparable to mAβ, indicating partial degradation, in agreement with the fluorescence analysis. However, by 24 h, β-sheet levels increased again, approaching those to fAβ levels, suggesting potential exhaustion of the degradation observed at 12 h (Supplementary Fig. 3E).

Unordered β-sheets increased before declining at 24 h (Supplementary Fig. 3F), aligning with the transition to structured β-sheets. Antiparallel β-sheets followed a similar trend, remaining high early but decreasing over time, consistent with fibril formation (Supplementary Fig. 3G).

These findings collectively highlight a complex non-linear dynamics for Aβ aggregation within microglial cells. Our *in vitro* experimental design represents a pathological condition where microglial cells are overloaded with an excess of fibrillar Aβ resembling the plaque microenvironment. Our data challenges the assumption that microglia constantly clear Aβ and highlights a dual role in both degradation and aggregation.

### Galectin-3 enhances microglial uptake and intracellular processing of Aβ in a conformation-dependent manner

To investigate how Gal3 regulates microglial handling of Aβ, we used murine microglial BV2 cell line with a permanent deletion of Gal3 (51) and supplemented recombinant Gal3 extracellularly. Aβ dynamics were monitored by live-cell imaging using the Operetta CLS System and fluorescent monomeric Aβ (mAβ) and fibrillar Aβ (fAβ) species (Fig. 4A, B, E).

**Figure 4.**
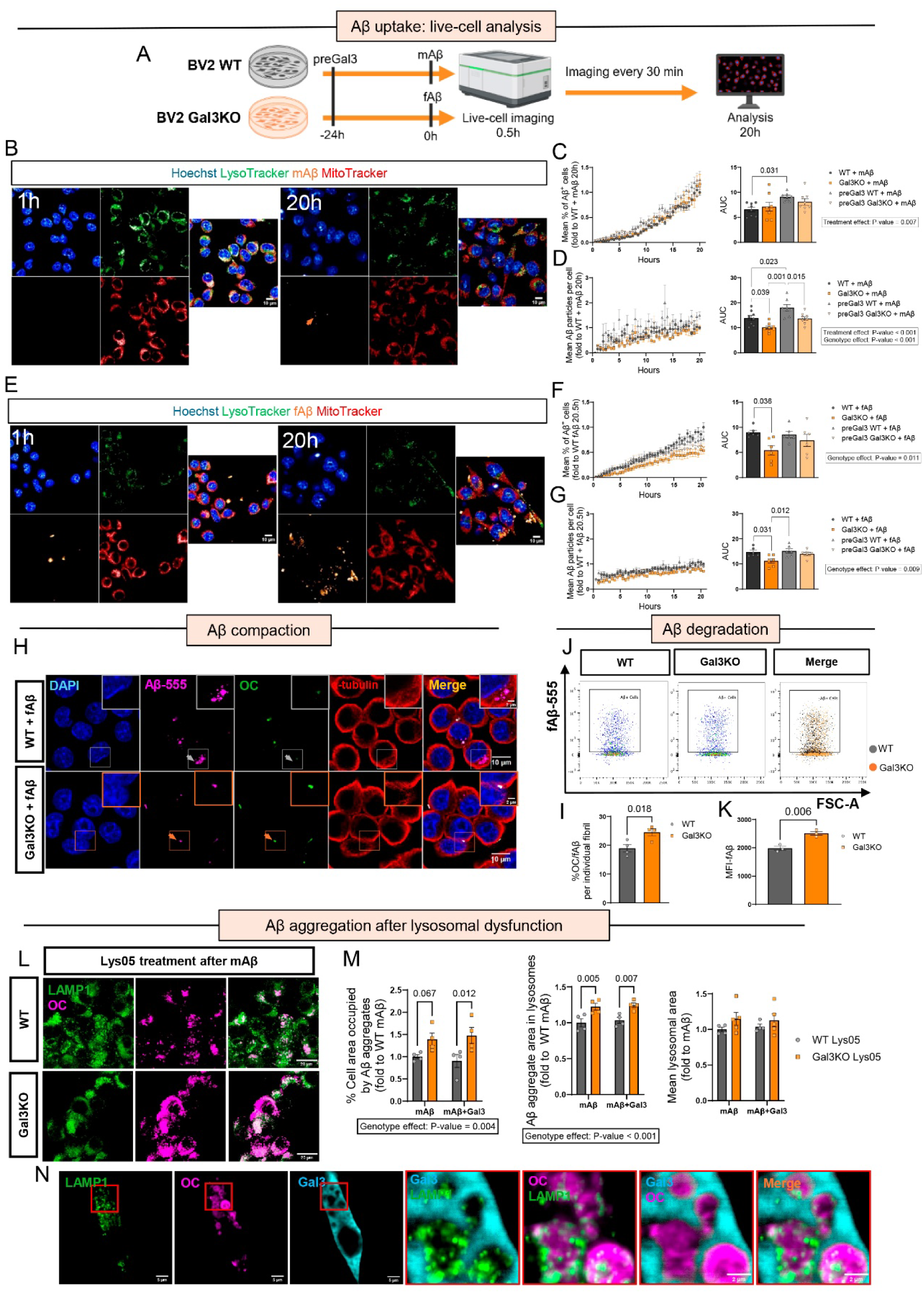
Galectin-3 differentially regulates microglial uptake, degradation, and compaction of monomeric and fibrillar Aβ species. **A** Schematic experimental outline of BV2 live-cell imaging Aβ uptake experiment. **B** Live-cell imaging snapshots of BV2 WT cells at 1h (left) and 20h (right) of mAβ (orange arrow). **C, D** Analysis derived from panel A live-cell imaging measurements: proportion of cells positive for intracellular Aβ after adding mAβ (C) and number of intracellular Aβ particles after adding mAβ (D) with area under the curve (AUC) quantification (n≥7/group). **E** Live-cell imaging snapshots of BV2 WT cells at 1h (left) and 20h (right) of fAβ. **F, G** Analysis derived from panel C live-cell imaging measurements: proportion of cells positive for intracellular Aβ after adding fAβ (F) and number of intracellular Aβ particles after adding fAβ (G) with area under the curve (AUC) quantification (n≥6/group). **H** Immunofluorescence staining of Aβ (MOAB2) and anti-fibrillar Aβ (OC) after fAβ addition for 6h. Gray arrows represent lack of overlap between OC and fAβ in WT cells, while orange arrows indicate this overlap in Gal3KO cells. **I** Quantification of percentage of OC-positive fAβ structures (n=4/group). **J** Representative flow cytometry gating of Aβ+ cells in WT and Gal3KO cells after fAβ addition. **K** Median Fluorescence Intensity (MFI) of fAβ after 3h of degradation, assessed by flow cytometry (n=3/group). **L** Representative LAMP1 and OC immunostaining following Lys05 treatment in BV2 WT and Gal3KO cells exposed to mAβ. **M** Quantification of intracellular Aβ content and lysosomal area after Lys05 treatment. **N** Representative confocal imaging of Gal3 staining surrounding OC+ structures after Lys05 treatment. Values are expressed as individual experimental replicates with mean ±SEM. In C, D, F and G, two-way ANOVA with Tukey’s multiple comparisons was performed (Fisher’s LSD test was performed in M due to having only 4 families of comparisons). In I and K, unpaired t-test was performed. P-values are expressed with 3 decimals.

Gal3KO BV2 microglial cells, showed altered Aβ dynamics when compared with WT counterparts. While both genotypes showed initial responses to mAβ (Fig 4B,C) Gal3KO cells displayed reduced uptake over time (Fig. 4D). In the case of fAβ, Gal3KO cells showed both a lower proportion of Aβ^+^ cells and fewer Aβ particles per cell (Fig. 4F, G), indicating impaired internalization of fibrillar species.

Importantly, exogenous Gal3 enhanced mAβ uptake in WT cells, and rescued the uptake deficit in Gal3KO cells exposed to fAβ (Fig. 4D, F, G), suggesting that extracellular and intracellular Gal3 differentially regulate Aβ internalization depending on its aggregation state.

We next examined the intracellular fate of Aβ. Staining with OC antibody, which recognizes amyloid fibrils and fibrillar oligomers, revealed that although WT cells contained overall higher levels of internalized fAβ, these particles exhibited reduced OC-reactivity (gray arrows) compared to Gal3KO cells (orange arrows, Fig. 4H,I), indicating lower fibrillar compaction.

Given that lysosomal dysfunction may fundamentally impair cellular degradative capacity, we sought to evaluate whether Gal3 influences the efficiency of Aβ degradation. First, cells were exposed to fAβ, washed, and incubated in serum-free medium prior to analysis by flow cytometry. Gal3KO cells exhibited increased Aβ fluorescence compared to WT cells, indicating a reduced degradation of internalized aggregates (Fig. 4J,K and Supplementary Fig. 4C). This phenotype was further supported by live-cell imaging, which showed in Gal3KO cells an increase in the intensity and size of large surface-associated Aβ aggregates (Supplementary Fig. 4B, D). No toxicity was observed under these conditions (Supplementary Fig. 4E).

Together, these findings indicate that Gal3 regulates microglial Aβ uptake and intracellular processing in a conformation-dependent manner. While Gal3 promotes Aβ internalization, it effectively limits the accumulation of aggregated species by supporting lysosomal degradative capacity. In contrast, Gal3 deficiency leads to differential Aβ processing and increased retention of fibrillar Aβ.

### Galectin-3 modulates microglial lysosomal responses to amyloid stress

Given the prominent colocalization of Gal3 within LAMP1^+^ lysosomal compartments within the microglia of AD brains, we next examined how Gal3 deletion affects lysosomal function under amyloid stress. To this end, we monitored lysosomes dynamics in BV2 cells using LysoTracker, marker of acidic vesicles during exposure to different Aβ species.

Gal3KO cells showed a modest reduction in basal lysosomal acidification, which became more pronounced following fAβ exposure, particularly after 12 hours (Supplementary Fig. 5A). This temporal shift coincided with the structural changes in the Aβ detected by s-μFTIR, suggesting a transition from effective processing to lysosomal overload.

To further characterize lysosomal function prior to exhaustion, we performed immunostaining for LAMP1 and Cathepsin B after 6 hours of mAβ or fAβ exposure. LAMP1 is a marker for endolysosomal vesicle abundance, and Cathepsin B, a luminal lysosomal protease involved in Aβ degradation. Similarly to Lysotracker, Gal3KO cells showed a modest reduction in basal LAMP1 intensity regardless of the Aβ treatment (Supplementary Fig. 5B). Also, Gal3KO cells exhibited reduced Cathepsin B levels compared to WT (Supplementary Fig. 5B). Importantly, colocalization analyses revealed decreased recruitment of Cathepsin B to LAMP1^+^ compartments in Gal3KO cells, and an overall increase after fAβ treatment, the later indicating no lysosomal membrane rupture caused by fAβ (Supplementary Fig. 5C). Overall, these points to a less degradative lysosomal capacity in Gal3KO cells providing a mechanistic basis for the increased fibrillar Aβ species observed in Gal3KO (Fig. 4I,K).

To directly assess the impact of lysosomal impairment on Aβ handling, we pharmacologically inhibited lysosomal acidification using Lys05. BV2 WT and Gal3KO cells were first exposed to mAβ for 5 hours, followed by Lys05 treatment for 72 hours in the presence or absence of exogenous Gal3, and subsequently analyzed by immunostaining for LAMP1 and OC. In accordance with previous research (52), Lys05 treatment increased both the intracellular content of OC^+^ aggregates and lysosomal area (Supplementary Fig. 5D, E). Notably, Gal3KO cells exhibited a more pronounced increase in both the amount and size of aggregated Aβ following Lys05 treatment, compared to WT cells (Fig. 4L, M), supporting a role for endogenous Gal3 in maintaining lysosomal-dependent processing of Aβ. In contrast, the addition of exogenous Gal3 did not alter Lys05-induced Aβ accumulation in either genotype, indicating that this effect is primarily mediated by intracellular rather than extracellular Gal3.

### Galectin-3 modulates Aβ aggregation dynamics by inhibiting secondary nucleation and stabilizing intermediate species associated with neurotoxicity

Having established that Gal3 regulates microglial processing of Aβ, we next examined whether Gal3 directly interacts with Aβ and alters its aggregation dynamics. Given that Gal3 is secreted under conditions of cellular stress, and present in the extracellular plaque environment, we tested its effect on Aβ fibrillization in a cell-free system.

Aggregation of mAβ was monitored using Thioflavin T (ThT) fluorescence in the presence of increasing concentrations of recombinant Gal3 (Fig. 5A). Gal3 delayed fibril formation in a dose-dependent manner, indicating an inhibitory effect on Aβ aggregation kinetics. Kinetic modelling using AmyloFit (53) revealed that Gal3 selectively impaired secondary nucleation, with minimal effect on primary nucleation or elongation (Fig. 5A, Supplementary Fig. 6A), suggesting a specific role in limiting aggregate proliferation (54). Consistent with this, Native PAGE analysis revealed altered aggregation patterns in the presence of Gal3, including the appearance of ∼60 kDa species indicative of intermediate or stabilized assemblies (Fig. 5B), while SDS-PAGE showed a rapid reduction in monomeric Aβ (∼5 kDa) (Fig. 5C), supporting a shift toward non-fibrillar intermediate species. Structural analysies using negative-stain transmission electron microscopy (TEM) and cryo-EM further demonstrated that Gal3 alters fibril morphology, resulting in structurally distinct fibrils (Fig. 5D, E).

**Figure 5.**
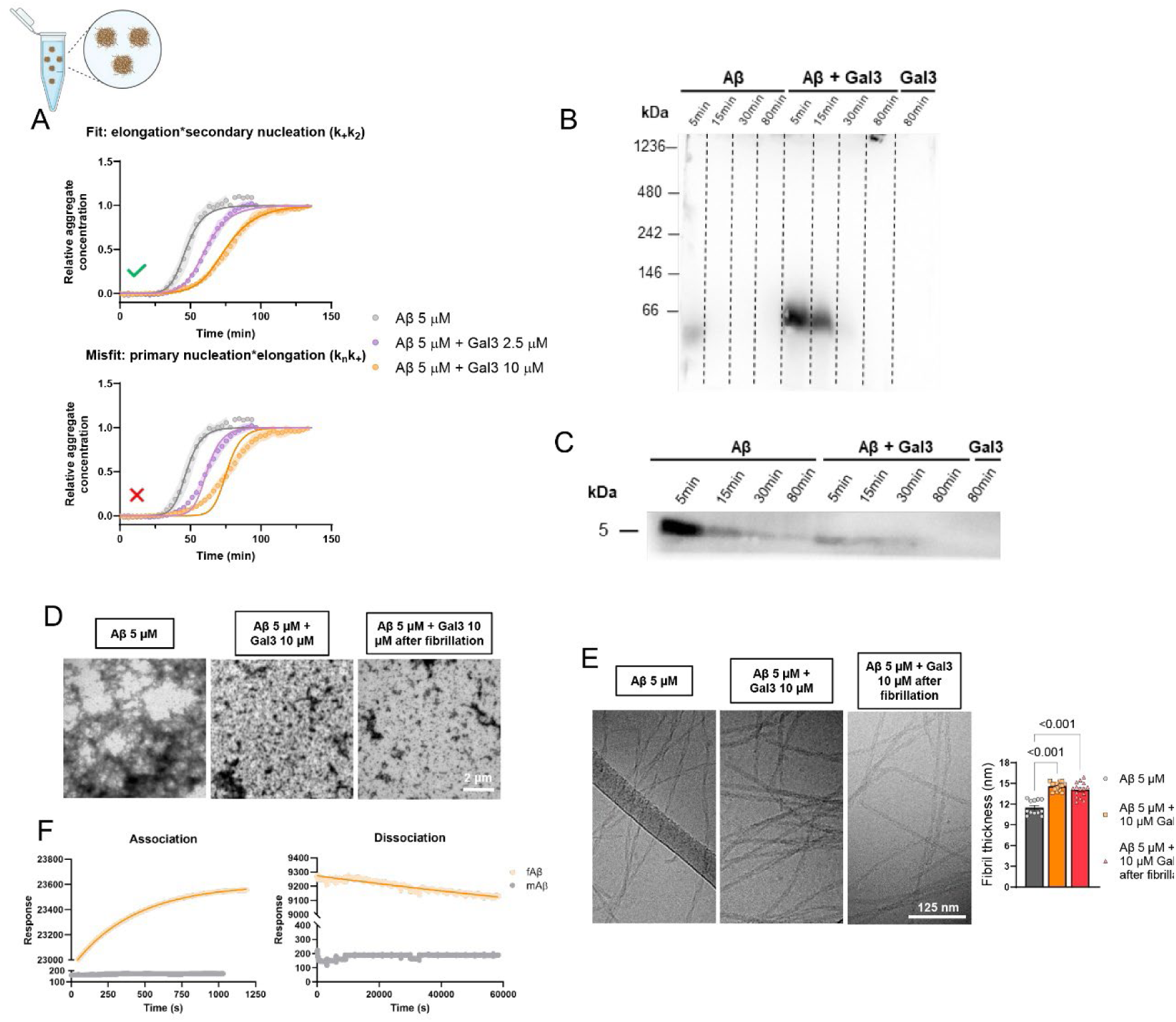
Extracellular Galectin-3 disrupts Aβ fibrillization by inhibiting secondary nucleation and binding fibrillar intermediates. **A** Ab42 fibril formation monitored as a function of time through Thioflavin-T (ThT) fluorescence intensity in the presence of varying concentrations Gal3 (n=8/group). The kinetic traces were alternately fitted using the secondary nucleation-dominated model of AmyloFit [Meisl, 2016 #269], allowing the product of k_+_k_2_ to vary freely (elongation and secondary nucleation, left) and k_n_k_+_ (primary nucleation and elongation, right), while the other product was kept as global constant at the value obtained for Aβ alone. The best fit was obtained for k_+_k_2_, represented with a green tick (left). **B** Native PAGE Western Blot anti-Ab (6E10) of different Ab fibrillations over time with and without Gal3 protein. Timepoints were selected based on a ThT assay which was run before starting the incubations (Supplementary Fig. 7B). **C** SDS-PAGE Western blot anti-Ab (6E10) of different Ab fibrillations over time with and without Gal3 protein. Same timepoints as in Fig. 5C were applied (based on Supplementary Fig. 7B). **D** Representative TEM images of Ab after fibrillation with and without Gal3 (first two images) plus Gal3 added only after fibrillation (third image). **E** Representative Cryo-EM images and quantification of fibril thickness (n≥12/group) of Ab after fibrillation with and without Gal3 (first two images) plus Gal3 added only after fibrillation (third image). **F** SPR sensograms, response as a function of time, reporting on the interaction of Gal3 with immobilized mAb (gray), and fAb (orange). Solid lines show fits to the data during association (left), and dissociation (right). The data has been background-subtracted and is represented as the average of two measurements. Time zero is set at the beginning of the injection (left) and after the injection (right). In A and F, values are expressed as mean ±SD. In E, each value represents an individual fibril thickness and mean ±SEM is represented. One-way ANOVA with Tukey’s multiple comparisons was performed and p-values are shown.

To determine whether these effects were mediated by direct interaction, we performed surface plasmon resonance (SPR) and found that Gal3 binds Aβ with high affinity, showing a marked preference for fibrillar over monomeric species (Kd of 9 ± 3 nM) (Fig. 5F).

Together, these findings indicate that Gal3 directly interacts with Aβ and modulates its aggregation dynamics by limiting secondary nucleation and stabilizing low-molecular weight species. These effects are consistent with the reduced Aβ compaction observed *in vitro* and *in vivo* and support a model in which Gal3 promotes the persistence of structurally distinct assemblies.

### Gal3 deletion is associated with altered microglial immune receptor programs and reduced neuroinflammatory signatures

To identify potential Gal-3-associated microglial receptors, we performed *in silico* structural screening AlphaFold-based protein-protein interaction prediction. Screening of 24902 human proteins identified 650 high-confidence interactors (pDockQ > 0.4). Notably, these candidates were enriched for leucine-rich repeats and immunoglobulin-like domains (Fig. 6A), structural motifs characteristic of key immune receptors, including TLR4 and TREM2 (55). Among these, TLR4 ranked among the top predicted interactors (21st), while modeling suggesting binding of Gal3 to its extracellular domain (Fig. 6B). Consistent with these predictions, TLR4 levels were reduced in Gal3KO BV2 cells upon fAβ exposure (Supplementary Fig. 2H, I), in agreement with previous evidence showing that Gal3 interacts with TLR4 to promote microglial activation (15). Moreover, absence of Gal3 promoted a more protective microglial profile, potentially involving TREM2 upregulation (Supplementary Fig. 2G, J). Functional enrichment of predicted Gal3 interactors using STRING revealed significant association with immune-related pathways, including TNF and chemokines (Fig. 6C, D), further supporting a role for Gal3 in modulating inflammatory responses. To determine the functional consequences of Gal3 deletion *in vivo*, we performed RNA-seq on FACS-sorted microglia from 18-months-old APP and APP-Gal3KO mice (Fig. 6E). Differential expression analysis revealed significant downregulation of multiple AD-associated genes (e.g., *Rpph1*, *Pla2g4b*, *Nedd4*, *Septin3*, *Tnr*, *Oas2*, *Dab1*), alongside decreased expression of the immunoreceptor *Cd300lf* (Fig. 6F). In contrast, genes associated with neuroprotection and inflammatory resolution, including *Rnase4* and *Tnfaip3,* were upregulated in APP-Gal3KO microglia (56, 57), suggesting a shift toward a less inflammatory transcriptional profile (For a full list of gene expression results, see Supplementary Table 3).

**Figure 6.**
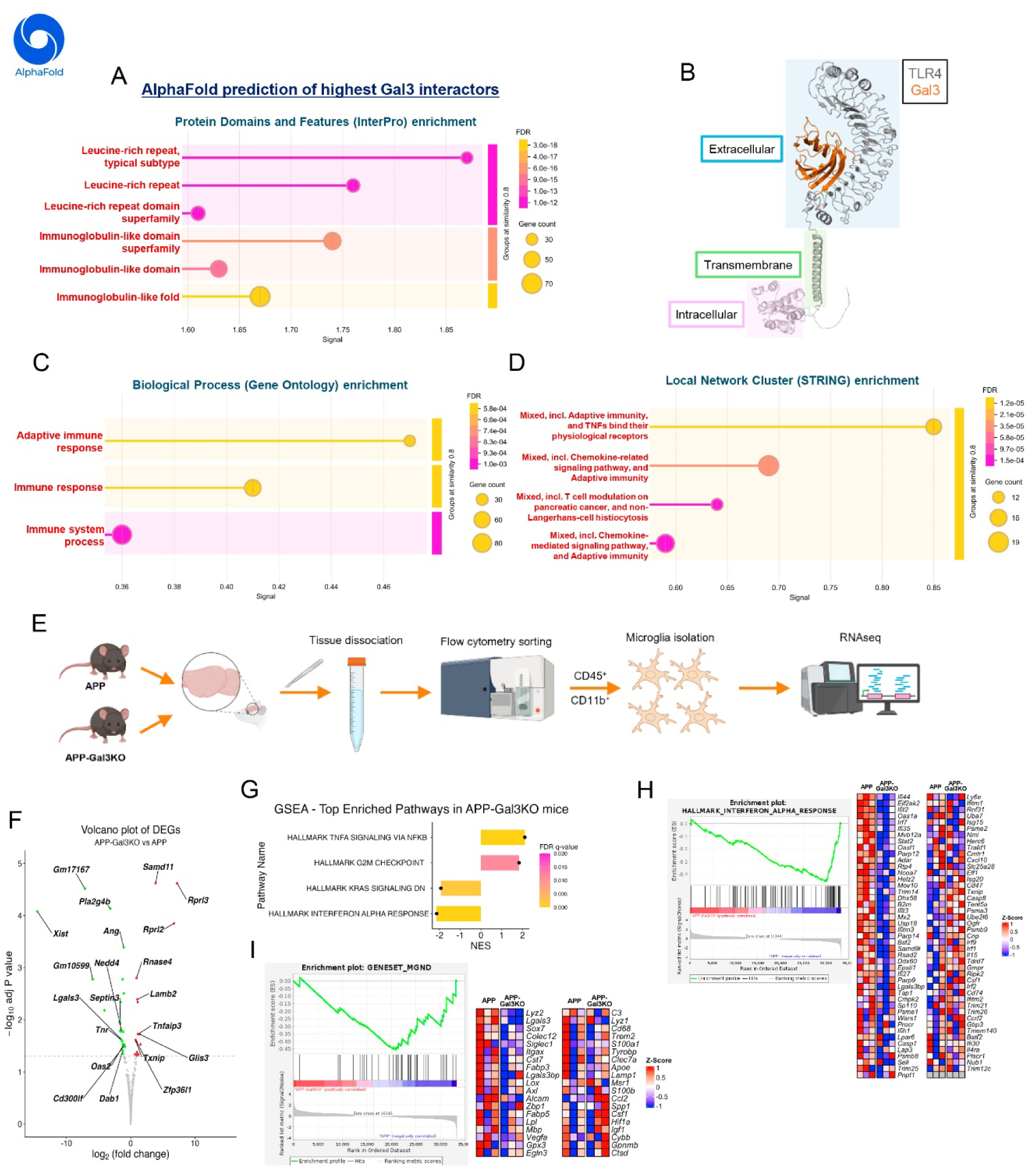
Galectin-3 interacts with microglial immune receptors and drives neuroinflammatory transcriptional reprogramming in AD. **A** Identification of enriched domains in the proteins predicted to interact with Gal3 (pDockQ score>0.4, using AlphaFold2), using STRING. For full list of Gal3 interactors, see Supplementary Table 2. **B** Structural model of Gal3 (highlighted in orange) in complex with TLR4 isoform D (highlighted in grey) generated using PyMOL. pDockQ score = 0.626 (Rank 21 in Fig. 5A). Spiral structures represent α-helix and parallel pointed structures represent β-sheet. **C** Identification of enriched biological processes in Gal3 interactors, using STRING. **D** Identification of enriched Local Network Clusters in Gal3 interactors, using STRING. **E** Experimental outline of RNAseq from isolated microglia. **F** Top Differentially Expressed Genes (DEGs) in APP-Gal3KO microglia compared to APP microglia. Rpph1 not shown in graph. **G** Top enriched pathways in APP-Gal3KO microglia compared to APP microglia using GSEA ranked by q-value and Normalized Enrichment Score (NES). **H** GSEA enrichment plot and heatmap using interferon-alpha response gene-set. I GSEA enrichment plot and heatmap using neurodegenerative microglia (MGnD) response gene-set.

Gene set enrichment analysis (GSEA) revealed reduced type I interferon (IFN-α) signaling and altered TNF-α/NF-κB pathway in APP-Gal3KO microglia (Fig. 6G, H), accompanied by decreased expression of IFN-stimulated genes such as *Oas2*. Consistently, the MGnD (microglial neurodegenerative) signature (4) was significantly negatively enriched in APP-Gal3KO cells (p = 0.0022, Fig. 6I), further supporting a shift away from the disease-associated microglial states (Full enrichment results of IFNα and MGnD in Supplementary Table 4). Upstream regulator analysis using Ingenuity Pathway Analysis (IPA) further supported a reduction in inflammatory signaling pathways along with activation of protective TNF pathways (Supplementary Fig. 6C, D; full list in Supplementary Table 5).

Together, these findings indicate that Gal3 deletion reconfigures microglial receptor interaction and signaling, suppressing detrimental neuroinflammatory programs.

## Discussion

Our findings identify Gal3 as a central regulator of microglial responses to Aβ, linking Aβ aggregation dynamics, lysosomal function, and neurotoxicity across human tissue, mouse models, and *in vitro* systems.

We show that Gal3⁺ microglia are selectively associated with cored plaques in AD patients, a plaque subtype strongly linked to neuritic dystrophy and neurotoxicity (34, 35). The presence of Gal3 in both microglial lysosomal compartments, dystrophic neurites and extracellular amyloid structures positions it at the interface between Aβ processing and extracellular plaque remodeling. Our data further demonstrate that extracellular Gal3 directly modulates Aβ aggregation dynamics. *In vitro*, Gal3 delays fibril formation by selectively inhibiting secondary nucleation, and rapidly destabilizing monomeric Aβ in favor of intermediate Aβ species. These intermediate assemblies are widely associated with increased neurotoxicity (47–50, 58), suggesting that Gal3 does not inhibit aggregation, but rather redirects Aβ toward structurally distinct and potentially more toxic states. This mechanism is consistent with the reduced plaque compaction observed *in vivo* and in human tissue, where Gal3⁺ plaques display larger halos and increased microglial engagement, and Gal3⁺ structures are only found in the periphery of the plaque core.

Our findings fit well with our previous work in Parkinson’s disease (29), where Gal3 was shown to directly interact with α-synuclein and remodel fibrillar assemblies into shorter, structurally altered strains with increased neurotoxic potential. In that context, Gal3 did not prevent aggregation but instead disrupted fibril integrity and promoted the formation of smaller, more reactive species associated with enhanced cellular toxicity. Remarkably, genetic deletion of Gal3 led to increased accumulation of α-synuclein aggregates but reduced neurodegeneration, highlighting a similar functional paradox. Together, these findings suggest that Gal3 may represent a conserved modulator of amyloid aggregation across neurodegenerative diseases.

At the intracellular level, our results identify the lysosomal compartment as a critical site where Gal3 regulates Aβ fate. Gal3 localized prominently to LAMP1⁺ compartments (20, 59), while lysosomal accumulation has been shown to accompany microglial activation in 5xFAD mice (60). Indeed, Gal3 deletion affected lysosomal homeostasis leading to decreased acidity and reduced Cathepsin B, suggesting impaired degradative capacity. Notably, lysosomal deacidification experiments revealed that microglia can generate fibrillar Aβ from monomeric species under conditions of lysosomal dysfunction. These findings suggest that dysfunctional lysosomes may act not only as sites of impaired degradation but also as intracellular environments that enable the conversion of monomeric Aβ into fibrillar aggregates. This effect is exacerbated in Gal3-deficient cells, highlighting the importance of intracellular Gal3 in regulating lysosomal-dependent Aβ processing under stress conditions. Notably, homeostatic microglia have been reported to initiate amyloid plaque seeding (28). In this context, our data support a model in which Gal3-expressing microglia actively contribute to amyloid remodeling, influencing the structural evolution of Aβ and potentially shaping the downstream pathological outcome.

Despite this role in promoting lysosomal processing, our data reveal a functional paradox. Gal3 enhances Aβ uptake, yet its presence is associated with increased neuritic dystrophy, reduced plaque compaction, and a more neurotoxic microenvironment. In contrast, Gal3 deletion resulted in more compact plaques, reduced dystrophic neurites, and increased TREM2 coverage, a phenotype associated with improved amyloid containment and reduced toxicity. Thus, TREM2 signaling promotes microglial clustering and plaque compaction while limiting Aβ diffusion and neurotoxicity (27, 30, 31, 38, 39). This observation suggests that sustained microglial uptake of Aβ precipitates lysosomal dysfunction, while the direct interaction between Aβ and Gal3 promotes the formation of proteotoxic species. Ultimately, this molecular cascade may drive aberrant microglial activation, shifting the cellular response from a protective state toward a chronic, neurodegenerative phenotype. These findings are consistent with previous studies linking microglial activation states to neurodegenerative phenotypes, including MGnD and type I interferon-driven responses (4, 6, 8, 9). Indeed, our transcriptomic and *in silico* analyses indicate that Gal3 shapes microglial inflammatory signaling by interacting with immune receptor domains characteristic of TLR4 and TREM2, and its deletion reprograms microglia toward a less inflammatory and less disease-associated state, with reduced type I interferon signaling and negative enrichment of the MGnD signature. This shift, together with increased TREM2 coverage *in vivo*, suggests that Gal3 modulates the balance between detrimental and protective microglial activation states. This effect is not unique to Gal3, as Gal9 also promotes microglial activation and exacerbates Aβ pathology in AD (61, 62). In contrast, astrocyte-derived Gal1 deactivates microglia and prevents CNS inflammation (63). Further studies should demonstrate whether different glyco-immune checkpoints act at different stages of microglia activation, differentiation and function to positively or negatively control neuroinflammatory and neurodegenerative responses (64).At the same time, persistent lysosomal engagement may drive intracellular stress and contribute to the formation of dystrophic neurites. In this context, Gal3 expression may become maladaptive for microglia: although it enhances Aβ uptake and partially preserves lysosomal function, it also promotes chronic remodeling of Aβ into structurally harmful species, a process that may extend beyond microglia through Gal3 secretion into extra-microglial sites such as dystrophic neurites. Rather than acting solely as a mediator of Aβ clearance, Gal3 reshapes amyloid pathology by regulating Aβ aggregation, intracellular processing and inflammatory signaling. This integrated role positions Gal3 as a key driver of stage-specific, neurotoxic microglial response in AD. Notably, Gal3-driven modulation of microglial function has also been observed in other neuropathologic conditions including models of ischemic injury (65), cuprizone-driven demyelination (66) and Parkinson’s disease (29).

From a therapeutic perspective, these findings suggest that targeting Gal3 may shift Aβ aggregation and microglial activation toward less toxic states. Ongoing clinical efforts targeting Gal3, including monoclonal antibodies (Phase 1b/2a, NCT05074498) and BBB-permeable small-molecule inhibitors (67, 68), highlight the importance of further understanding Gal3 function in AD pathogenesis.

In conclusion, our data identify Gal3 as a key link between lysosomal dysfunction, Aβ aggregation remodeling, and neuroinflammatory signaling. Through these interconnected processes, Gal3 promotes the development of a neuroinflammatory and neurotoxic microenvironment in AD, highlighting its critical role as a promising therapeutic target.

## Materials and Methods

### Additional material and methods, as well as full antibody list available in Supplementary Information

#### Human samples

Hippocampal tissue from AD cases (*n* = 9) (The Netherlands Brain Bank) were analyzed in the study. (subject information is provided in Supplementary Table 1). Written informed consent for the use of brain tissue and clinical data for research purposes was obtained from all patients or their next of kin following the International Declaration of Helsinki and Europe’s Code of Conduct for Brain Banking. The medical ethics committee of VU medical center in Amsterdam approved the procedures for brain tissue collection, and the ethical review authority board in Sweden approved the study.

#### Immunofluorescence staining of human samples

At autopsy, the hippocampal samples were formalin-fixed (for approximately 15 h), immersed in 30% glucose for 2-3 days, and thereafter sectioned into 40-µm-thick sections, which were kept free-floating in cryoprotectant solution at -20°C. For immunofluorescence staining, sections from (n = 9) AD cases were rinsed (3 × 10 min) in 0.02 mol/L potassium phosphate-buffered saline (KPBS). Next, the sections were blocked in blocking solution (KPBS with 0.25% Triton X-100, Sigma-Aldrich, Cat: T8787) and 5% normal donkey serum) for 1 hour on a shaker at room temperature. Thereafter, the sections were incubated with primary antibodies diluted in blocking solution overnight on a shaker at 4°C. Primary antibody used: Iba1 (1:500 dilution), MOAB2 (1:500 dilution), Gal3 (1:750 dilution), LAMP1 (1:200 dilution). The following day, the sections were rinsed (6 × 10 min) in KPBS with 0.25% Triton X-100 and thereafter incubated with the corresponding secondary antibodies diluted in blocking solution. Finally, the sections were incubated with Methoxy-04 in KPBS (1:1000 dilution from 10 mg/mL, Bio-techne, Cat: 4920) for 1 h at room temperature (RT). After rinsing (3 × 10 min) in KPBS, the sections were mounted on slides and left to dry in darkness. Thereafter, the mounted sections were incubated for 5 min in 1% Sudan Black (Sigma-Aldrich) dissolved in 70% ethanol. Subsequently, after being rinsed in KPBS, mounting medium and coverslips were added. Stained tissues were imaged with an Olympus VS120 Slide Scanner and a Nikon A1RHD confocal microscope. For super-resolution microscopy images, Nikon N-SIM E microscope was used.

#### Animals

The hAPP751sl (+/-) murine model, referred to in this text as APP, is a transgenic model that overexpresses a mutated form of the human amyloid precursor protein (isoform 751). Specifically, this protein carries both the Swedish mutation (K670N/M647L) and the London mutation (V717I) under the control of the murine Thy1 promoter (Sanofi). APP mice were crossed with *Lgals3^-/-^*(Gal3KO) mice (Dr K. Sävman, Gothenburg University, Sweden) to obtain APP/*Lgals3^-/-^* model. All animal models share a common C57BL/6 wild type background. They were kept at constant humidity (60%) and temperature (22°C), on 12 h dark / 12 h light cycles and access to food and water ad libitum. Experiments were carried out in accordance with the Guidelines of the European Union Directive (2010/63/EU) and Spanish regulations (RD53/2013 and BOE 34/11370-421, 2013) for the use of laboratory animals; the study was approved by the Scientific Committee of the University of Seville.

APP/PS1 mice usage for immunoelectron microscopy experiments was carried out in accordance with the Spanish and the European Union regulations (RD53/2013 and 2010/63/UE), and with the approval of the committee of Animal Research from the University of Malaga (Spain).

#### Immunofluorescence

For immunofluorescence staining, mice were anesthetized with isoflurane and transcardially perfused with saline buffer (0,9% NaCl in water). Brains were extracted and saved for 7 days on 4% PFA at 4°C. Lately, they were stored at least 48 h on a 30% sucrose 0,1% sodium azide solution until they were frozen in dry ice for 15 minutes. Following the freezing procedure, brains were serially sectioned at 30 μm thickness slices in the coronal plane on a Leica CM1850 cryostat. The slices were stored at -40°C on antifreeze solution (30% sucrose, 30% ethylene glycol in phosphate buffer 0,1M pH 7,4).

After slicing, free floating sections from APP and APP-*Lgals3^-/-^* were first rinsed 3 times with PBS for 5 minutes per rinse. In the case of staining ThS, BAM10, TREM2 and LAMP1, tissues were incubated at 80°C on citrate buffer for 30 min. Next, tissues were permeabilized by their incubation with PBS-Triton X100 1% (PBS-T) for 1 hour and blocked with 2 h of PBS-T 1% with 5% bovine serum albumin (BSA) for 2 h. After blocking, tissues were incubated with primary antibodies, diluted in PBS-T 0,1% BSA 1%, for 30 min at room temperature and, lately, over 18 hours at 4°C. Primary antibody used: BAM10 (1:500), LAMP1 (1:200), TREM2 (1:200), CD68 (1:750). Following this incubation, tissues are placed at room temperature for 30 min. 6 rinses with PBS-T 0.1% of 5 min each are carried out before an 1 h incubation with the corresponding secondary antibodies diluted in PBS-T 0.1% BSA 1%. For ThS (1:1000 dilution from 10 mg/mL, Sigma-Aldrich, Cat: T1892) or DAPI staining, its addition takes place after secondary antibodies incubation for 5 min, diluted in PBS-T 0,1%. Next, 3 rinses in PBS-T 0,1% and other 3 rinses in PBS are performed. Finally, brain sections were mounted in glycerol 50% on slides. Slides and coverslips were sealed with nail polish. Visualization was performed on a Leica Stellaris 8 confocal microscope.

#### Cell lines

Murine microglial BV2 cell lines (RRID: CVCL_0182) were cultured in Dulbecco’s Modified Eagle Medium GlutaMax (DMEM GlutaMax, Gibco, Cat: 61965059) supplemented with 10% heat inactivated Fetal Bovine Serum (FBS, Gibco, Cat: 10270106) and 1% penicillin-streptomycin (Cytiva, Cat: SV30010) and kept at 37°C 5% CO_2_. Gal3-deficient BV2 Gal3KO cells were generated as previously described (51).

#### Live-cell imaging experiments

For live-cell imaging experiments, 6000 cells were seeded on a 96-well ViewPlate microplate (Revvity, Cat: 6005225. After cellular attachment to the plate bottom, 1 μM of Gal3 protein was added 24 h prior to initiating imaging (preGal3). The following intracellular compartments were labeled through the addition of cell-penetrating dyes: mitochondria (using MitoTracker Deep Red FM at 1:2500 dilution, Invitrogen, Cat: M22426), lysosomes (using LysoTracker Green DND-26 at 1:2500 dilution, Invitrogen, Cat: L7526) and nuclei (using Hoechst 33342 at 1:1000 dilution). Lastly, 1 μM mAβ-555, 1 μM fAβ-555 or sterile PBS (Gibco, Cat: 10010056) was added. 8 images per well were taken every 30 min for 20 h with Operetta CLS High Content Screening (PerkinElmer) by confocal spinning disk technology with a 40× water immersion objective (experiments with fAβ were imaged for 0.5 h longer because of system availability reasons).

#### Lys05 experiment

10.000 WT and Gal3KO cells were seeded on 96-well ViewPlate microplates under standard culture conditions as previously described. After 5 h, the medium was replaced with medium containing 0.5% FBS and Poly(I:C) (polyinosinic:polycytidylic acid, 10 μg/mL, Sigma Aldrich, P0913) to induce microglial activation, and cells were maintained overnight (14 h). Cells were then treated with mAβ (0.25 μM) for 5 h. Subsequently, medium containing Lys05 (0.5 μM, Selleck Chemicals, S8369) and Gal3 (1 μM) was added. After 72 h, cells were fixed with 4% PFA for subsequent immunocytofluorescence analysis.

#### Immunocytofluorescence

6000 cells were seeded on a 96-well ViewPlate microplate. After cellular attachment to the plate bottom overnight, they were treated with fAβ-555 for 6 h. They were then fixed with 4% PFA and washed with PBS. They were permeabilized with PBS and 0.1% Tween-20 (PBS-T20) for 10 min and blocked with PBS-T20 with 10% Normal Donkey Serum (PBS-T20-NDS) for 1 h. Then they were incubated overnight at 4°C with PBS-T20-NDS using the following primary antibodies: anti-amyloid fibrils OC (1:1000 dilution), anti-cathepsin B (1:500 dilution), anti-LAMP1 (1:100 dilution) and anti-α-tubulin (1:1000 dilution). After washing with PBS, cells were incubated with appropriate secondary antibodies in PBS 10% NDS in darkness. All secondary antibodies were Alexa Fluor IgG (1:500 dilution). After washing with PBS, cells were stained with DAPI solution (1:1000 dilution, Thermo Scientific, Cat: 62248) for 5 min.

A Leica SP8 with 63× oil objective using laser scanning confocal microscope was employed to image samples. Multiple images in a z-stack with a 2 μm step size were taken in mouse brain samples.

#### Thioflavin-T assay and fitting

The thioflavin-T (ThT) fluorescence intensity was measured using FLUOstar Omega plate readers (BMG LABTECH) in PEGylated polystyrene (non-binding surface) 96-well plates. Fibril formation was monitored via the ThT fluorescence intensity at 480 nm using an excitation wavelength of 448 nm measured through the bottom of the wells at 37 °C.

The kinetic traces were fitted using the online fitting platform AmyloFit (53) using either the unseeded secondary nucleation model or the seeded secondary nucleation model, with the initial seed concentration set to zero. Kinetic traces for 5 mM Ab alone were fitted and it was tested which of the obtained rate constants could be used as global constants and which needed to be locally fitted variables to get reasonable fits to the kinetic data obtained in the presence of Gal3.

#### Transmission Electron Microscopy

To study the aggregated Aβ, 400 mesh copper grids were coated with Pioloform and afterwards with carbon. Later, the grids were treated in a glow discharge equipment. fAβ was then added and after letting them dry, 2% uranyl acetate was used as a negative stain. fAβ were examined in a transmission electron microscope (TEM, Tecnai BioTwin 120 kV), equipped with an Olympus Veleta camera.

#### Cryo-EM

Lacey formvar/carbon 200 mesh copper grids (Ted Pella) were glow discharged in a Gloqube (Quorum). The grids were prepared using a Leica EM GP where sample was applied, blotted and plunge-frozen in liquid ethane. Imaging of the grids was done in a JEM-2200FS (Jeol) with an Omega filter and TVIPS F416 camera.

#### STRING analysis

The list of potential partners of Gal3 was analyzed using the STRING network database (69) available here: https://version-12-0.string-db.org/cgi/network?networkId=bhnwDjNZpEwY). 466 nodes were analyzed by the database.

#### Microglial isolation

For microglia isolation, the procedure begins with animal perfusion and the extraction of the cerebral cortex. The tissue is then mechanically dissociated using a scalpel and passed through 20G needles in HBSS 1X with 10% FBS. Next, the sample is filtered through a 70 µm sieve pre-moistened with dissociation buffer and centrifuged at 160 g for 10 min at 4°C with the brake applied. The pellet is then resuspended in an isotonic Percoll solution (90% Percoll + 10% HBSS 1X) at 30%, and a gradient is created with HBSS 1X. It is centrifuged again at 800 g for 10 min at 4°C with the brake applied. The supernatant is discarded, and the pellet is washed with PBS 1X. After another centrifugation at 800 g for 10 min at 4°C with the brake applied, the pellet is resuspended in 500 µL of PBS 1X. Subsequently, 400 µL of the sample is mixed with 1 µL of anti-CD11b-APC antibody and 1 µL of anti-CD45-PE antibody. This mixture is incubated for 20 min at room temperature in the dark. Finally, the samples are processed using the FACS Aria Fusion cell sorter (Becton Dickinson).

#### Bulk RNA Sequencing

RNA quality was examined using the Bioanalyzer® 2100 (dsDNA HS kit) to inspect DNA size, and quantified by Qubit assay. cDNA libraries were prepared using SMART-Seq Stranded Kit (Takara, Cat: 634444). Barcoded adapters were added to allow combining of libraries for sequencing. Size distribution was assessed with Bioanalyzer DNA High-Sensitivity Chip and quantified with Qubit fluorometer. Libraries were pooled in equimolar concentrations and sequenced on an Illumina NextSeq 500 platform (Mid Output flow cell) using 2×75 bp paired-end chemistry. On average, 25-27 million read pairs were obtained per sample.

#### Statistical analyses

All statistical analyses were performed using GraphPad Prism 10 (GraphPad). Statistical tests used, as well as value of N can be found in figure legends, and exact p-values are indicated in each graph.

#### Figure creation

Schematic figures were partly generated using biorender.com (under publication license).

## Acknowledgments

We thank Pontus Nordenfelt and Vibha Kumra Ahnlide (Lund University) for the training and usage of N-SIM microscope. Sara Linse (Lund University) for the purification and labelling of Aβ42-S8C and for input in SPR analysis. We also thank the Cell and Gene Therapy Core (Lund Stem Cell Centre) for the design of the CRISPR- KO experiment. We thank the Lund Protein Production Platform (LP3, www.lu.se/lp3; Lund University) for the production of recombinant human Gal3. Lund University Bioimaging Centre (LBIC), Lund University, is gratefully acknowledged for providing experimental resources. LU-Fold protein structure prediction facility has to be thanked for the AlphaFold2 predictions. These computations were enabled by the supercomputing resources Berzelius provided by the National Supercomputer Centre (NSC) at Linköping University and the Knut and Alice Wallenberg foundation. Additional computational resources used were provided by the National Academic Infrastructure for Supercomputing in Sweden (NAISS) and the Swedish National Infrastructure for Computing at NSC, Chalmers University Centre for Computational Science and Engineering (C3SE), and PDC Centre for High Performance Computing, KTH Royal Institute of Technology, partially funded by the Swedish Research Council through grants 2018-05973 and 2022-06725). Thanks to Crispin Hetherington for the expert help with Cryo-EM imaging. This work was supported by the Strategic Research Area MultiPark at Lund University, Lund, Sweden (2020-2025); the Swedish Research Council (2018-03033); the Swedish Alzheimer Foundation (AF-9685, 2021); Olle Engkvist Foundation (188-0100, 219-0166), Swedish Brain Foundation (21-0387, 2021); the A.E. Berger Foundation (F210040, 2021); G&J Kock Foundation (2020) to T.D.; the Spanish Ministerio de Ciencia e Innovación/ Agencia Española de Investigación (PID2024-157400OB-I00)) to J.L.V. The Spanish Ministerio de Ciencia e Innovación/ Agencia Española de Investigación (PID2023-146447OA-I00), The Royal Fysiografen Foundation (45376 and 44192 to A.B.-S) and by grants PI21/00915 and PI24/00274 from Instituto de Salud Carlos III Spain (cofinanced by FEDER funds from the European Union) to J.V and A.G., respectively. G.A.R. is supported by the Minister of Science and Technology, Fundación Sales and Fundación Bunge & Born (Argentina) as well the CaixaResearch Institute (Spain).

