## Supplementary Information for "Galectin-3 drives neuroinflammatory amyloid remodeling by stabilizing intermediate Aβ species and altering lysosomal processing"

**This PDF file includes:**

SI Methods

Supplementary Figures 1 to 6

Supplementary Table 1

Captions for Supplementary Tables 2 to 5

Antibody list

SI References

**Supplementary information methods**

**Protein production**

*Galectin-3*

Recombinant human Gal3 was produced by the Lund Protein Production Platform (LP3). Gal3 production were prepared as previously described (1), and and no detectable endotoxin contamination was found using the Pierce™ Chromogenic Endotoxin Quant Kit (Thermo Scientific). Freeze-dried Gal3 was stored at -80°C until use.

*Recombinant amyloid-beta*

Prior research indicates that synthetic Aβ has failed to induce a robust activation of cultured innate immune cells (2) and the effect of incorporating a fluorophore into the protein structure has been shown to affect its aggregation process (3). For this reason, in-house isolation was performed as follows: amyloid beta 42 wild-type with an N-terminal Met, Aβ(M1-42), (MDAEFRHDSGYEVHHQKLVFFAEDVGSNKGAIIGLMVGGVVIA), and the S8C mutant, Aβ(M1-42)S8C, MDAEFRHDCGYEVHHQKLVFFAEDVGSNKGAIIGLMVGGVVIA, were expressed in *E coli* and purified as previously described (4, 5). To ensure ultra-pure and homogeneous starting material, fresh Aβ monomers (mAβ) were purified just prior to the kinetics, surface plasmon resonance and cryo-EM experiments. This was done by reconstituting an aliquot of freeze-dried Aβ42 in 6 M guanidine hydrochloride followed by size exclusion chromatography (SEC) on a Superdex 75 Increase 10/300 (GE Healthcare) column equilibrated in the desired buffer with a flow rate of 0.5 mL/min on a Bio-Rad NGC system. All buffers were degassed and filtered with wwPTFE 0.2μM 50 MM disc filters (Pall corporation). mAβ concentrations were determined by integrating the chromatograms in ChromLab (BioRad) using an extinction coefficient of 0.31 ((mg/ml)^-1^cm^-1^) and molecular mass of 4.645 kDa. For cell treatment, the freshly isolated mAβ were aliquoted and flash-frozen in liquid nitrogen before being aliquoted and stored at -80°C and thawed in wet ice for experiments. Each aliquot was thawed only once, but this freeze-thaw cycle could impair the complete monomeric status of the protein. During purification of Aβ(M1-42)S8C, 1 mM DTT (PanReac Applichem, Cat: 3483-12-3) was included in all buffers to ensure monomeric peptide, except in the final monomer isolation step to produce peptide for fluorophore labelling with Alexa Fluor 555 using maleimide chemistry to label the unstructured N-terminal region as previously described (6). Monomer (mAβ-555) was isolated twice using SEC as above to remove any traces of free dye. The tagged version of Aβ contains a cysteine amino acid in the eighth residue (instead of a serine) to attach the Alexa fluorophore covalently, this labelling position has been found to minimally retard the aggregation process if used at as a minor component in mixtures with unlabeled peptide with no observable effect on the resulting fibril structure (6).

For thioflavin-T (ThT) fluorescence, surface plasmon resonance, and cryo-EM experiments, 20 mM sodium phosphate, 0.2 mM EDTA pH 8.0 was used. In the first case, 10 μM ThT was included, and for SPR, 0.005% Tween-20 (T20, Sigma-Aldrich, Cat: P1379) was included. mAβ for cell studies was isolated the same way but without EDTA and subjected to sterile filtering by passing the monomer through a 0.2 μm syringe filter (Sarstedt) prewashed in deionized water and 1 mL 5 μM mAβ to saturate the filter to ensure no adsorption or loss when sterile filtering the subsequent monomer.

To achieve fibrillar forms of the peptides, solution starting from mAβ and mAβ-555, were incubated overnight at 50 µM at 37°C in Protein LoBind tubes (Eppendorf), obtaining fAβ and fAβ-555 respectively. Labelled peptide was always used at a 10:1 ratio of WT:Ab42S8C-Alexa555.

**Synchrotron-Based FTIR Microspectroscopy**

Synchrotron-based Fourier-transform infrared (s-FTIR) microspectroscopy (µFTIR) was conducted at the SMIS beamline of the SOLEIL synchrotron (L’Orme des Merisiers, 91192 Gif-sur-Yvette Cedex, France). BV2 cells were seeded on a sterilized 1-mm calcium fluoride (CaF₂) coverslip (Crystran Ltd, Cat: CAFP10-1) and allowed to adhere. 30.000 cells were added on the coverslips, and were treated with 5 µM fAβ for either 1 h, 12 h or 24 h; and with 5 µM mAβ for 24 h. Then, cells were washed with PBS and fixed with 4% PFA for 15 min followed by 2 washes with PBS and one wash with 20 mM HEPES. Finally, coverslips were left to air dry.

The single-point infrared measurements were performed in transmission mode using a ThermoFisher Scientific Continuum XL FTIR microscope equipped with a 32× magnification, 0.65 NA Schwarzschild objective. Spectra were recorded over a spectral range of 4000-1000 cm⁻¹ at a resolution of 4 cm⁻¹, with a 6 µm × 6 µm aperture and 256 co-added scans. Single-point spectra were acquired for each cell.

Acquired spectra were analyzed using Quasar software (University of Ljubljana, Slovenia). Quality assessment and noise artifact identification, including Mie scattering effects, were performed using k-means clustering and principal component analysis (PCA). Preprocessing steps included spectral normalization, baseline correction, and second derivative transformation using a Savitzky-Golay filter (7-point window, third-order polynomial) to enhance spectral differentiation, particularly in the Amide I region.

MATLAB 2024a (MathWorks) was used to visualize the spectra.

**Cytotoxicity experiment**

20.000 WT and Gal3KO cells were plated on 96-well plates (Sarstedt). The next day, mAβ or fAβ were added at different concentrations (0.5 μM, 1 μM, 3 μM and 5 μM). After 24 h, the cell medium was analyzed using CyQUANT™ LDH Cytotoxicity Assay (Invitrogen, Cat: C20300) following manufacturer’s instructions.

**Flow cytometry analysis**

For degradation experiments, 50.000 cells were seeded on 48-well plates and treated with mAβ-555 or fAβ-555 for 1 h in starvation medium. After that, medium was removed and cells were washed once with PBS, and fresh medium with reduced serum was added. At different timepoints (0, 3, 24, and 48 h) cells were detached and analyzed as following: they were detached by removing medium, washing once with sterile PBS and adding trypsin 2.5% (Gibco, Cat: 15-090-046) for 2 min. Cells were then spined down in fresh medium and resuspended in FACS buffer (PBS+2% FBS). Fc-receptors were blocked using CD16/CD32. Afterward, they were stained with CD45-AlexaFluor 488 (1:200 dilution) and CD11b-PE-Cy7 (1:200). PI (1:1000 dilution) was used as viability dye for cells. Cells were analyzed in BD FACSAria III Cell Sorter. Data were analyzed with FlowJo software version 10.4.0.

**Surface plasmon resonance data analysis**

The dissociation of Gal3 from fAβ was first fitted to the blank-subtracted sensogram region after the end of the injection using Kaleidagraph and the equation: $y=A*e^{-koff*t}$. The best fit was obtained with a dissociation constant, $k_{off}$ = $4.3\cdot{10}^{-6}$. The association of Gal3 to fAβ was fitted to the sensogram during Gal3 injection. This part of the data was obtained as the section after the signal jump caused by the change in refractive index when the analyte is injected in the flow cell, until the end of the injection, and the time of the injection start is treated as time zero. The data were uploaded to the AmyloFit online platform (7) and fitted using the custom equation: $A+B\cdot m_{0}\cdot k_{on}\cdot\frac{1-\exp\left( -\left( m_{0}\cdot k_{on}+k_{off} \right)\cdot t \right)}{\left( m_{0}*k_{on}+k_{off} \right)}$ with $k_{off}$ kept constant at $4.3\cdot{10}^{-6}$, while the constant, A, max amplitude, B, and $k_{on}$ were varied in the fit.

**Surface plasmon resonance studies**

The association between mAβ or fAβ and Gal3 was investigated using surface plasmon resonance (SPR) technology with a BIAcore 3000 instrument. 20 mM sodium phosphate pH 8.0 with 0.005% T20 was used as running buffer for all SPR experiments.

fAβ were immobilized on a CM3 sensor chip by amine coupling with 37.5 g/L 1-ethyl-3-(3-dimethylaminopropyl) carbodiimide (EDC) and 6.75 g/L N-hydroxysuccinimide (NHS) in water injected at a flow rate of 10 μL/min on all four channels of a CM3 sensor chip to generate reactive succinimide esters. 20 μM sonicated fAβ in 20 mM sodium phosphate pH 8 were diluted to 1.0 μM in 10 mM sodium acetate pH 3.0 and injected in channels 3 and 4. 20 μM mAβ was diluted to 1.0 and 2.0 μM in 10 mM sodium acetate, pH 3 and injected in channel 2. 1 M ethanolamine-HCl, pH 8.5 was injected in all four channels to deactivate any remaining esters.

This yielded a CM3 chip with a ligand-free channel 1 as a control surface, monomer covalently coupled in channel 2 and fibril coupled in channel 3 and 4. All further injected species were diluted in the running buffer before being injected over the chip.

**Native PAGE Western Blot**

Electrophoresis under native conditions was performed in NuPAGE 3 to 8%, Tris-Acetate gels (Invitrogen, Cat: EA0375) and the following steps were performed as described in Klementieva et al. (8). Primary antibody used was anti-β-Amyloid 1-16 (6E10, 1:1000 dilution). Secondary antibody used was horse anti-mouse IgG peroxidase (1:3000 dilution).

**SDS-PAGE Western blot**

To study microglial markers, 50000 cells were seeded in 24-well plate. After 24 h, the medium was changed to starvation medium (medium with 2% FBS) and 1.0 μM of Gal3 (preGal3) was added to the cells. After 24 h, treatment with mAβ or fAβ was performed for 6 h. Then, the cell medium was centrifuged for 10 min at 10000 g 4°C and stored at -80°C. Cells were then treated with RIPA buffer (Fisher Scientific, Cat: 89900) with PhosStop and cOmplete phosphatase and protease inhibitor tablets (Roche, Cat: 4906847001 and Cat: 4693116001) on wet ice, extracts were sonicated for 10 sec, 80% amplitude, 0.8 cycles and stored at -80°C. After centrifugation for 10 min at 15000 g 4°C, supernatant was stored at -80°C.

For Gal3KO cell characterization, cells were extracted as described above.

Immunoblotting of proteins extracted from cell cultures or in the cell-free experiment was performed as previously described (9).

Primary antibodies used were: anti-galectin-3 (1:1000 dilution), anti-TREM-2 (1:1000 dilution), anti-TLR4 (1:1000 dilution), anti-β-Actin Peroxidase (1:10000 dilution) and anti-actin (1:1000 dilution). For the cell-free experiment, anti-β-Amyloid 1-16 6E10 (1:1000 dilution) was used.

Secondary antibodies used were: horse anti-goat IgG peroxidase, goat anti-rabbit IgG peroxidase. For the cell-free experiment, horse anti-mouse IgG peroxidase was used. All secondary antibodies were used at 1:3000.

Outliers were identified through the Grubbs’ method using Alpha=0.1 (10).

**Immunoelectron microscopy**

Immunogold procedure was performed as previously described (11). Briefly, after deep anesthesia with sodium pentobarbital (60 mg/kg), 6-month-old male APP/PS1 transgenic mice (Sanofi, delivered by Charles River), expressing human mutant APP751 with Swedish [KM670/671NL] and London [V717I] mutations and PS1(M146L)) (12); were transcardially perfused with 0.1 M phosphate-buffered saline pH 7.4 followed by 4% paraformaldehyde (PFA, Histolab, Cat: 96753) containing 75 mM lysine and 10 mM sodium metaperiodate in 0.1 M PB, pH 7.4. Brains were post-fixed overnight in the same fixative solution at 4°C and coronally sectioned (50 μm) on a vibratome (Leica VT1000S). Free-floating sections were then incubated in anti-Gal3 (1:5000 dilution) in a Phosphate Buffer Saline (PBS) 0.1 M/0.1% sodium azide/2% BSA-solution for 48 h at 22°C. The tissue-bound primary antibody was detected by the incubation with the corresponding 1.4 nm gold-conjugated secondary antibody (1:100 dilution) overnight at 22°C. After postfixation with 2% glutaraldehyde, the labeling was enhanced with the HQ Silver Kit (Nanoprobes, Cat: 2012), and gold toned. Finally, immunolabeled sections were fixed in 1% osmium tetroxide, block stained with uranyl acetate, dehydrated in acetone, and flat embedded in Araldite 502 (Electron Microscopy Sciences, Cat: 13900). Selected hippocampal areas were cut in ultrathin sections (70-80 nm) and examined with an electron microscope (JEOL JEM 1400). No specific labelling was observed in control sections that were processed as above but omitting the primary antibody.

**Gal3 exposure to Aβ species for native PAGE, SDS-PAGE and electron microscopy**

To select the timepoints for Aβ incubation with Gal3 for native PAGE and SDS-PAGE electrophoresis and subsequent Western blotting, fibril formation was monitored on an aliquot of freshly isolated 10 μM mAβ using ThT with and without 20 μM Gal3, as well as 20 μM Gal3 on its own. While ThT was running, a different aliquot of peptides was kept on ice for later incubations. After the ThT assay finalized, the timepoints of 5 min, 15 min, 30 min or 80 min were selected based on the following notions: before fibrillation, at t_½_ of Aβ alone, at t_½_ of Aβ with Gal3 and after fibrillation plateaued (red lines in Supplementary Fig. 6B). Then, Aβ was incubated with or without Gal3 for those times, at the same conditions than the ThT assay but without ThT (with the same concentration, plate type, temperature and plate movement), using the aliquot that was maintained on ice.

To find the relevant time for Gal3 addition to Aβ in TEM and cryo-EM, freshly isolated 5 μM mAβ were incubated in 96-well plates overnight, with or without 10 μM Gal3. Fibril formation was monitored using ThT fluorescence in a replicate sample. Gal3 was added to the formed fAβ when its corresponding replicate had reached a plateau in the ThT assay. The actual samples used for imaging contained no ThT.

The higher molarity in the native and SDS-PAGE experiments was chosen to ensure enough protein content on the gels.

**Image analysis**

Human immunostainings were analyzed in two ways. First, full section scans were used for plaque characterization, in which each brain region was selected and independently analyzed. For this analysis, plaques were qualitatively quantified depending on plaque morphology, using Methoxy-X04 and MOAB2 staining as basis. Four types of plaques were consistently found throughout patients and regions, and therefore quantified:

- Inert (Methoxy⁺/Aβ⁻): Plaques positive for Methoxy-X04 but lacking Aβ antibody signal, indicating highly compacted fibrils with limited epitope exposure. These are likely mature, stable plaques with minimal microglial interaction.
- Compact (Methoxy⁺/Aβ⁺ with complete colocalization): Plaques showing full overlap of Methoxy-X04 and Aβ signals, representing dense, well-packed fibrillar cores with accessible Aβ epitopes.
- Filamentous (Methoxy⁺/Aβ⁺ with diffuse filaments): Plaques with partial colocalization, displaying irregular, thread-like Aβ filaments extending beyond the compact core, indicative of ongoing aggregation and increased microglial activity.
- Cored (dense Methoxy⁺ core with Aβ⁺ halo): Plaques with a dense fibrillar core stained by Methoxy-X04 surrounded by a peripheral Aβ⁺ halo of less compact, possibly oligomeric Aβ, often associated with axonal damage and dystrophic neurites.

Each plaque was then classified based on the presence or absence of Gal3-positive microglia (Gal3⁺). For comparisons of plaque types by Gal3 positivity across brain regions, regions with no or only one Gal3⁺ plaque were excluded. This ensured analysis focused on areas where Gal3⁺ plaques were sufficiently represented to determine which plaque types were most commonly associated with Gal3. This analysis was done combining OlyVIA v4.1 (Olympus) to select brain areas and Fiji (Fiji by ImageJ, v1.54p, NIH) for quantification.

Second plaque analysis consisted of individually imaging a series of randomly selected Gal3⁺ plaques and Gal3^-^ plaques and quantifying plaque area, plaque circularity and microglial coverage using Fiji. 5-30 Gal3⁺ plaques per subject, and 10-50 Gal3^-^ plaques per subject were quantified. To quantify microglial uptake of methoxy-positive structures in human brains, three Gal3⁺ and 3 Gal3^-^ plaques per subject were quantified using Imaris Analysis Software v10.2 (Oxford Instruments). The percentage of material B (Methoxy-X04) above threshold colocalized with Iba1 was used for quantification.

For correlations of dystrophic neurites in human samples with Gal3 reactivity, individual plaque quantification was done with Fiji and included 6 patients, with 14-27 plaques per patient included. Microglial Gal3 was quantified by integrated density of Gal3 in Iba1^+^ structures. Dystrophic neurites are quantified by integrated density of LAMP1 or AT8 (pTau) in LAMP1^+^Iba1^-^ structures or pTau^+^Iba1^-^ structures. Dystrophy-associated Gal3 was quantified by integrated density of Gal3 in dystrophic neurites (LAMP1^+^Iba1^-^ structures). Plaque core intensity was quantified by mean gray value of Methoxy-X04. Imaris Analysis Software was used for 3D reconstructions.

Immunofluorescence of mouse brains was analyzed using the Z Project function of Fiji software. Once applied the projection, ThS signal was thresholded in order to create a region of interest, subsequently dilated and processed by watershed. This mask was exported to each individual channel image. By thresholding and measuring limited to threshold, Aβ, Iba1, CD68, TREM2 and LAMP1 signals were measured. For cortical and hippocampal individual plaque quantification by ThS and Aβ, 75 plaques per mouse were analyzed. For CD68 and TREM2 reactivity, 50 plaques per mouse were analyzed in the cortex, and 20 in the hippocampus. For LAMP1 immunoreactivity, 100 plaques per mouse were analyzed in the cortex and 75 in the hippocampus.

For live-cell and other experiments using Operetta CLS, images were analyzed employing the High Content Analysis System Harmony. The results shown are the mean of at least 6 imaged fields per well (with intraexperimental well duplicates). For the rest of image analysis experiments, Fiji by ImageJ was used.

Specifically, for OC immunofluorescence in the experiment without Lys05 exposure, images were taken using Leica SP8 with 63× oil objective using laser scanning confocal microscope, and Fiji software was used to quantify as follows: the fAβ channel was thresholded and “Watershed” was applied. Then, the function “Analyze Particles” was used and all individual fibrils saved in ROI manager. OC channel was also thresholded and the fAβ particles were used as a mask to quantify the amount of OC (% Area) covering each particle. The average of OC % Area of all the fibrils per image was saved. 3 images per experimental replicate were analyzed. For OC immunofluorescence in the experiment with Lys05 exposure, cells were stained with Phalloidin-iFluor 647 (Abcam, Cat: ab176759), while imaging and quantification was done using Operetta CLS.

For fibril thickness quantification in Cryo-EM, the average thickness throughout each single fibril was measured (3-4 values per fibril were analyzed), and 4 different images adding 12 fibrils per sample were studied.

To evaluate the colocalization between LAMP1 and Cathepsin B upon fAβ treatment, the Mean Gray Value of each channel was studied posterior to background subtraction. Subsequently, Coloc2 plugin was used and Mander’s Overlap Coefficient was reported for percentage of above-background pixels in LAMP1 that overlap with Cathepsin B after thresholding (Mander’s tM1).

Outliers were identified through the Grubbs’ method using Alpha=0.1 (10).

Western blot band quantification was carried out using Image Lab Software (Bio Rad).

**Protein complex prediction**

The human proteome (GRCh38) was downloaded from NCBI (<https://www.ncbi.nlm.nih.gov/projects/genome>). MMseqs2 (13) (Release 13-45111) (13) was used to cluster the human proteome sequences (130184 sequences) which resulted in a representative set of sequences (24902 sequences). AlphaFold2 (AF2) (14), as implemented in FoldDock (15) was used to co-fold Gal3 (*LGALS3*, UniProt ID: P17931) against the representative human protein sequences in pairwise combinations. Here, a truncated version of Gal3 sequence was used (P115 - I253) to co-fold against the representative human sequences. Gal3-containing heterodimeric complexes were ranked based on the pDockQ (predicted DockQ) score for the protein-protein interface (Python script available from https://gitlab.com/ElofssonLab/huintaf2/-/blob/main/bin/analysepairs.py, 10Å used as the interface distance cut-off). High-scoring acceptable models are usually considered at DockQ ≥ 0.23 (15), but this study was more conservative, only considering pDockQ score of ≥ 0.4 as indicators of interacting surfaces (16) (full list in Supplementary Table 2). AF2 predicted structures of Gal3 (NCBI Reference Sequence: BAA22164.1) and Toll-Like Receptor 4 (TLR4, NCBI Reference Sequence: NP_612567) were visualized with PyMOL (Schrödinger, LLC).

**Bulk RNA data analyses**

Raw sequencing reads were demultiplexed, quality-filtered, and adapter-trimmed, including the removal of poly-A/T sequences, using FASTQ Toolkit v1.0.0. Read alignment and transcript quantification were performed using the nf-core/rnaseq pipeline (17, 18) (v3.17.0) (19) STAR (v2.7.11b) was used for genome alignment, and Salmon v1.10.3 (20) was employed for transcript-level quantification against the GENCODE GRCm39 reference genome (21) and annotation. Differential gene expression analysis was conducted using the nf-core/differentialabundance pipeline (v1.5.0) (17), utilizing DESeq2 v1.38.0 (22) for statistical analysis. All nf-core pipelines were executed using default settings unless modifications were explicitly stated. Multiple testing correction was applied using the Benjamini-Hochberg method, with differentially expressed genes defined as those with an adjusted p-value < 0.05. Gene set enrichment analysis (23) (GSEA) was performed using the hallmark gene collection from the Molecular Signatures Database (MSigDB) (24), as well as a curated gene set of upregulated genes from microglial in neurodegenerative phenotype (MGnD) as identified by Krasemann et al (25). Data visualization was performed using ggplot2 v3.5.1 (26) in R v4.3.1 (R Core Team).

**Ingenuity Pathway Analysis**

Ingenuity pathway analysis (27) (IPA, QIAGEN Inc., https://digitalinsights.qiagen.com/IPA) was employed to investigate predicted cellular functions and upstream pathways of the RNAseq microglial data. A normalized p-value of 0.05 and a log2fold change above 0.25 and below -0.25 was used.

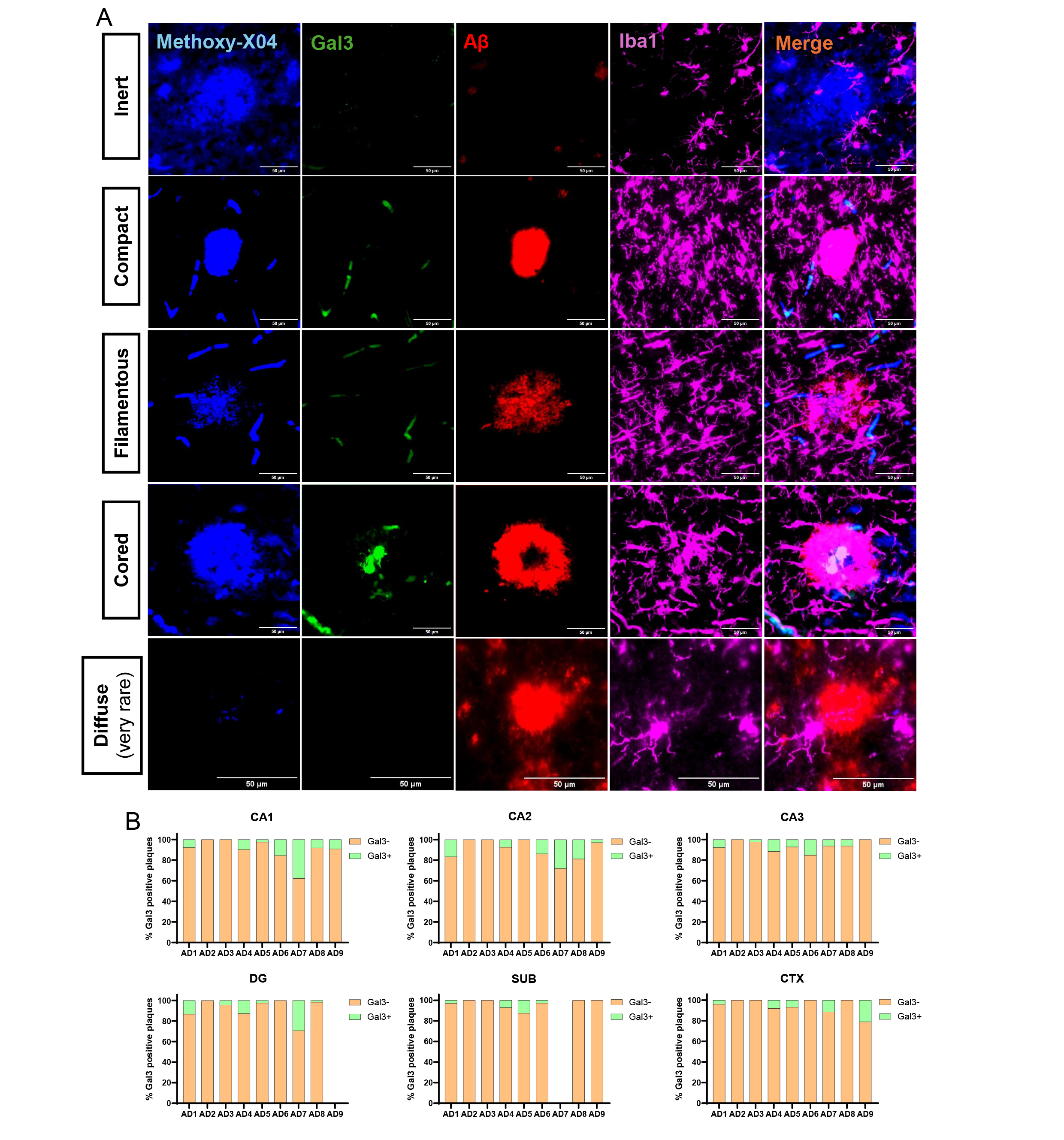

**Supplementary Figure 1**. **Postmortem human immunostaining plaque characterization. A** Representative human postmortem immunostaining showing the four plaque types analyzed plus the very rare diffuse plaque. **B** Individual subject proportions of Gal3^-^ and Gal3^+^ plaques across regions.

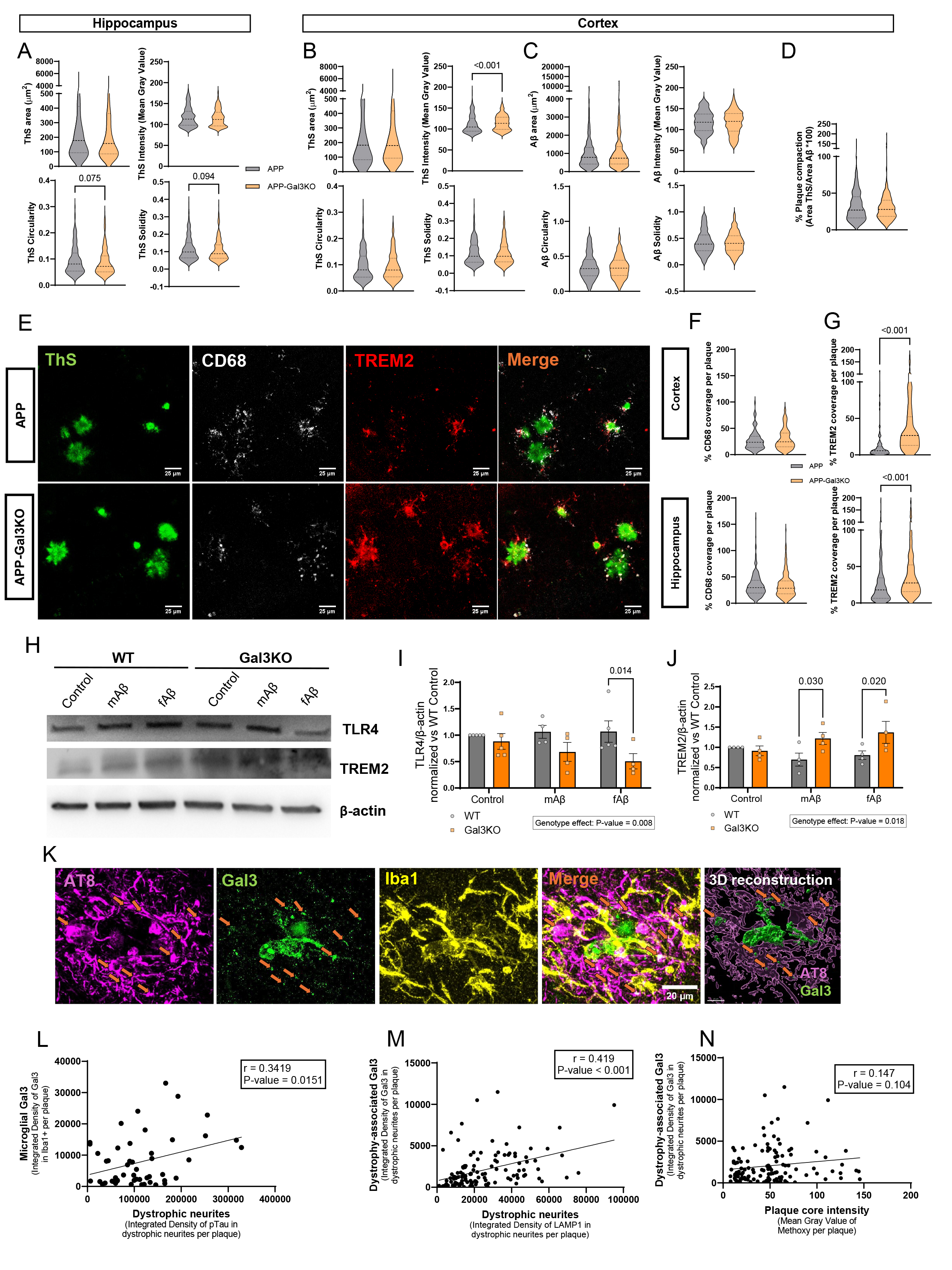

**Supplementary Figure. 2**. **Plaque morphology evaluation in APP and APP-Gal3KO cortex and human Gal3 relationship with dystrophic neurites. A, B** Morphology parameters of individual plaque cores by ThS quantified in hippocampus (A), cortex (B), n=75 plaques per mouse, ≥4 mice per group). **C** Morphology parameters of individual plaques by BAM10 (Aβ) quantified in cortex (n=75 plaques per mouse, ≥4 mice per group). **D** Proportion of plaque compaction measured by Area ThS/Area Aβ*100 in cortex and area covered by microglia per plaque measured by area of Iba1/Area Aβ*100 in cortex (n=75 plaques per mouse, ≥4 mice per group). **E** Representative individual plaque staining of microglial activation markers CD68 and TREM2, with ThS. **F, G** Proportion of plaque covered by CD68 (F) or TREM2 (G) in hippocampus and cortex (n=20 plaques per mouse, 5 mice/group). **H** Representative Western Blot images of TLR4 and TREM2 after the different Aβ treatments. β-actin was used as housekeeping. **I** Quantification of TLR4 Western Blot bands (n≥4/group). **J** Quantification of TREM2 Western Blot bands (n≥4/group). **K** Representative immunostaining and 3D reconstruction of dystrophic neurites, stained with pTau (AT8), Iba1 and Gal3. **L** Individual plaque correlation analysis between pTau^+^ dystrophic neurites density and microglial Gal3 density. **M** Individual plaque correlation analysis between dystrophic neurites density and Gal3 density in dystrophic neurites. **N** Individual plaque correlation analysis between plaque core intensity and Gal3 density in dystrophic neurites. In A, B, C, D, F and G; values are expressed as violin plots. In I and J, values are expressed as individual experimental replicates with mean ±SEM. In A, B, C, D, F and G; Mann-Whitney test was performed. In I and J, two-way ANOVA with Tukey’s multiple comparisons was performed. P-values are expressed with 3 decimals. In L, M, N, Pearson correlation was performed. P-values are expressed with 3 decimals.

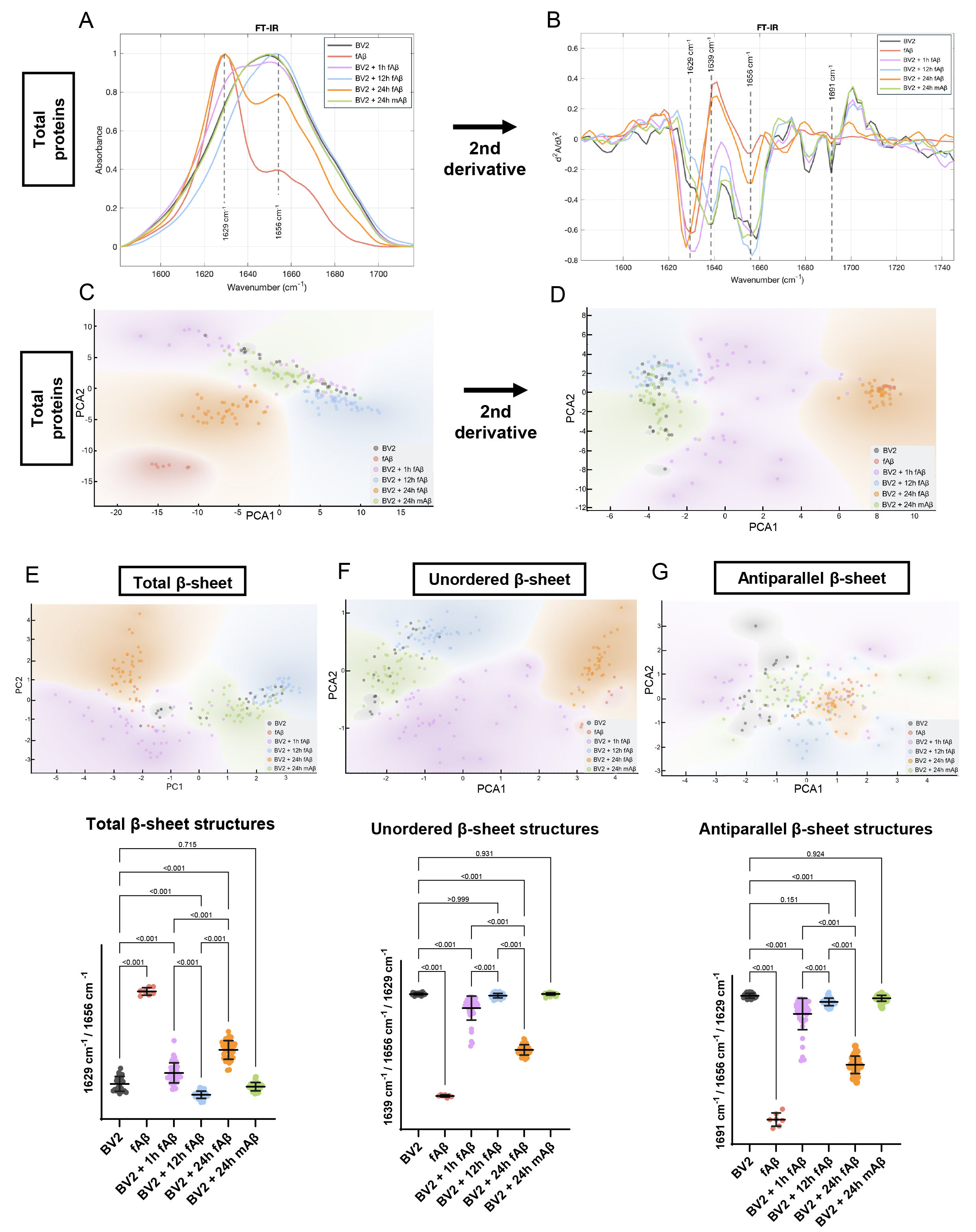

**Supplementary Figure 3.** **Microglial processing of Aβ aggregates involves dynamic structural remodeling revealed by synchrotron-based FTIR spectroscopy.** Key spectral features were quantified to assess β-sheet structure. **A** Preprocessed Fourier-transformed infrared (FT-IR) spectra. **B** Second derivative of FT-IR spectra. **C** Principal component analysis (PCA) of FT-IR spectra. **D** PCA of second derivative FT-IR spectra. **E** PCA and group comparison of the proportion β-sheet content by the intensity ratio of 1629 cm⁻¹ (β-sheet structures) to 1656 cm⁻¹ (total proteins). **F** PCA and group comparison of the proportion of unordered β-sheet content by the intensity ratio of 1639 cm⁻¹ (unordered β-sheet structures) to both 1629 cm⁻¹ (β-sheet structures) and 1656 cm⁻¹ (total proteins). **G** PCA and group comparison of the proportion of unordered β-sheet content by intensity ratio of 1691 cm⁻¹ (antiparallel β-sheet structures) to both 1629 cm⁻¹ (β-sheet structures) and 1656 cm⁻¹ (total proteins). In A and B, spectra were averaged for visualization. In C, D, E, F, and G, PCA was performed on spectral features associated with known structural components to identify clustering patterns. In E, F, and G, two-way ANOVA with Tukey’s multiple comparisons was performed. P-values of relevant comparisons are shown. In total, 38-45 treated cells per group were analyzed. For controls, 23 spectra were from untreated BV2 cells (negative control) and 7 spectra were collected directly from amyloid fibrils (positive control).

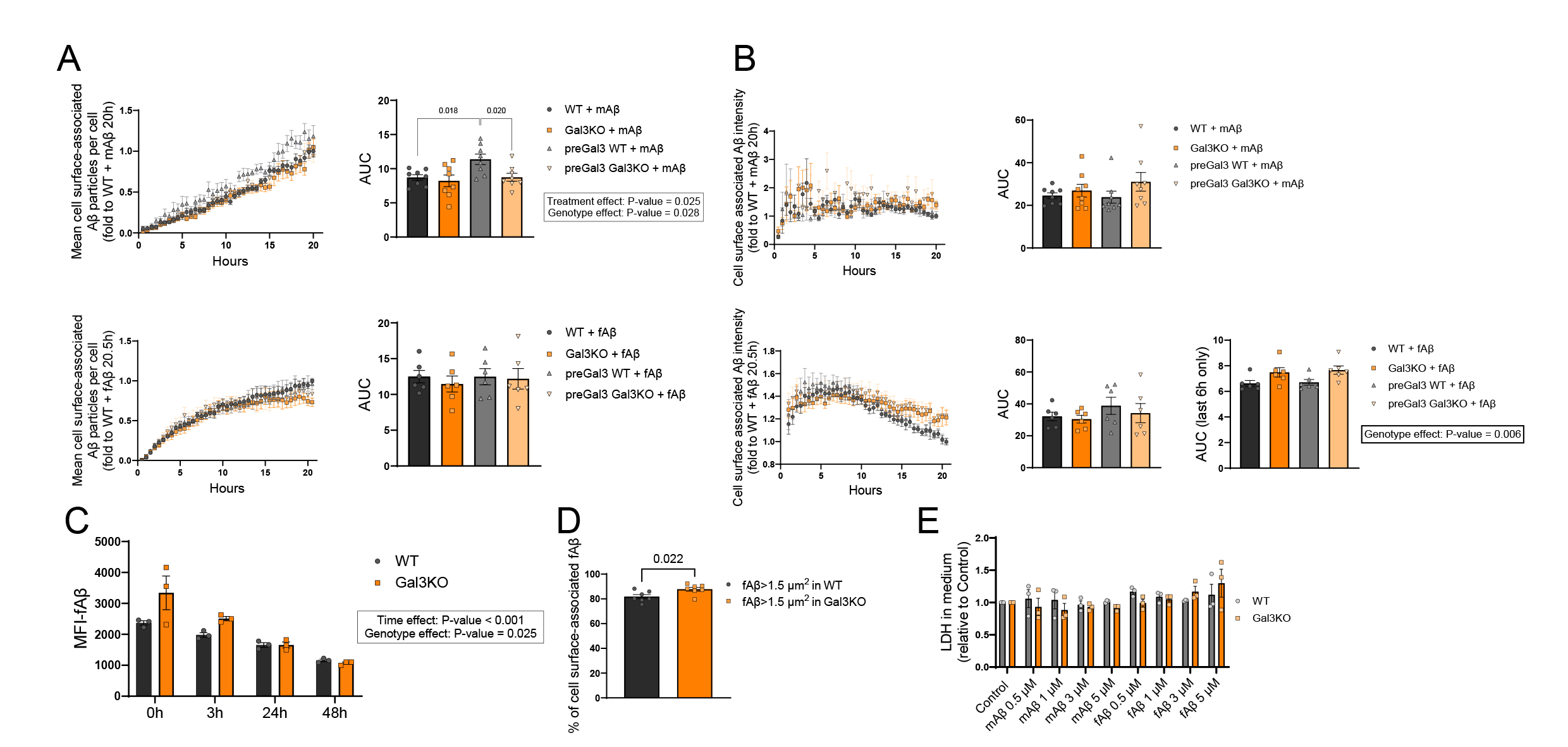

**Supplementary Figure 4. Number and area of Aβ particles attached to the cell membrane, intracellular and membrane-bound Aβ intensity, mitochondrial measurements and Aβ toxicity assay. A** Live-cell imaging data of number of Aβ particles attached to cell surface after adding mAβ (top) or fAβ (bottom), with area under the curve (AUC) quantification and analysis of results (n≥6/group). **B** Size of Aβ particles attached to cell surface after adding mAβ (top) or fAβ (bottom), with AUC quantification and analysis of results (n≥6/group). **C** Live-cell imaging data of intensity of intracellular Aβ particles after adding mAβ (top) or fAβ (bottom), with AUC quantification and analysis of results (n≥6/group). **D** Live-cell imaging data of intensity of Aβ particles attached to cell surface after adding mAβ (top) or fAβ (bottom), with AUC quantification and analysis of results (n≥6/group). **E** Number of mitochondrial particles after adding mAβ (top) or fAβ (bottom), with AUC quantification and analysis of results (n≥6/group). **F** Percentage of fAβ>1.5 μm^2^ attached to a cell surface respecting all fAβ in BV2 WT and BV2 Gal3KO (n=7/group). **G** Median fluorescence intensity (MFI) of fAβ in different degradation assays over 0, 3, 24 and 48 h (n=3/group). **H** Lactate dehydrogenase (LDH) levels in cell media after Aβ addition for 6h, relative to Control where no Aβ was added (n=3/group). Values are expressed as individual experimental replicates with mean ±SEM. Unpaired t-test was performed in F. Two-way ANOVA with Tukey’s multiple comparisons was performed in A, B, C, D, E and H. Two-way ANOVA without multiple comparisons was performed in G. P-values are expressed with 3 decimals.

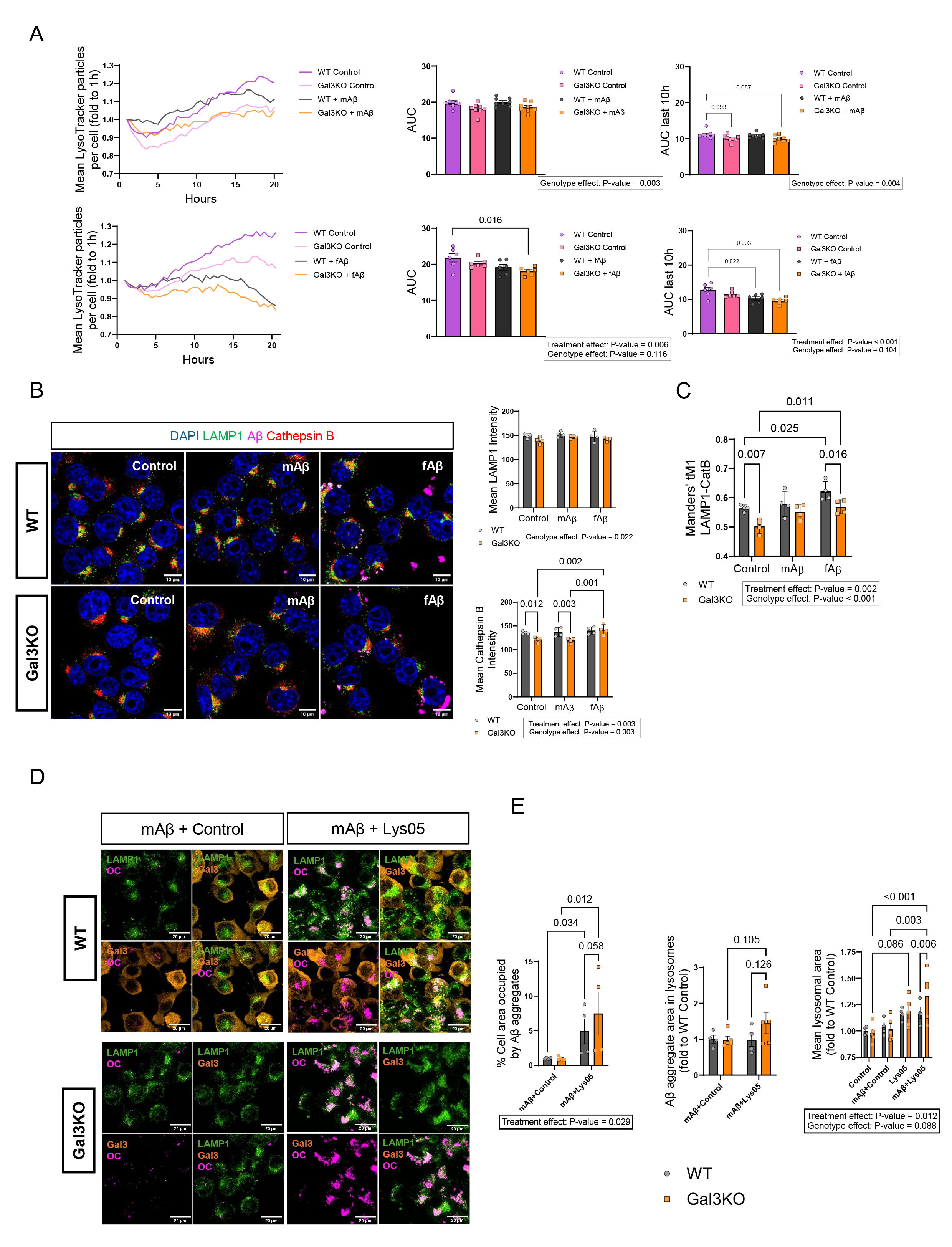

**Supplementary Figure 5**. **Galectin-3 deletion reshapes microglial lysosomal response to Aβ by limiting protease recruitment and vesicle acidification**. **A** Number of lysosome particles per cell after adding mAβ (top) or fAβ (bottom), with area under the curve (AUC) quantification and analysis of results (n≥6/group); with AUC re-analysis of the last 10 hours of the experiment. **B** Representative immunofluorescence staining of LAMP1 and Cathepsin B (CatB) and mean intensity quantifications after fAβ addition for 6 h (n=4/group). **C** Co-localization analysis of LAMP1 and CatB by thresholded Manders’ M1 (n=4/group). **D** Representative immunofluorescence staining of LAMP1 and anti-fibrillar Aβ (OC) in cells treated with mAβ with or without Lys05. In A, live-cell lines are expressed as mean values throughout replicates. The rest of values are expressed as individual experimental replicates with mean ±SEM. In A, B, C, two-way ANOVA with Tukey’s multiple comparisons was performed (Fisher’s LSD test was performed in E due to having only 4 families of comparisons). P-values are expressed with 3 decimals.

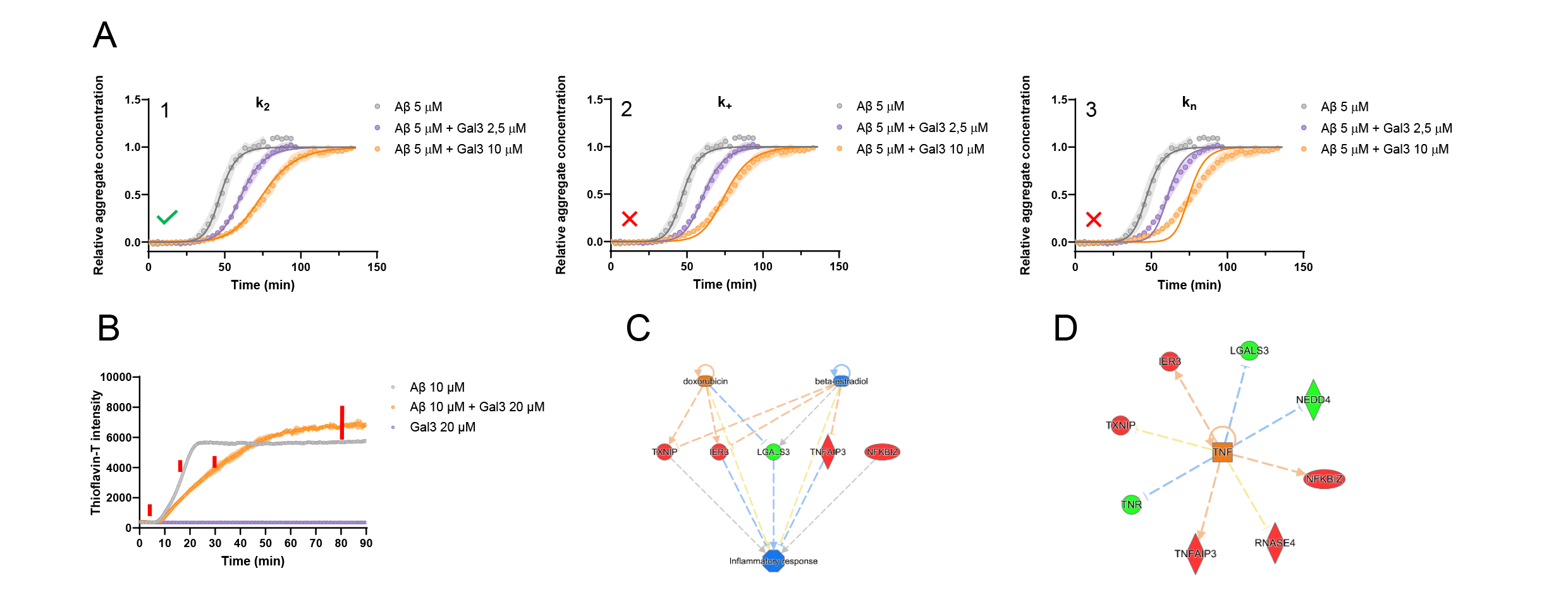

**Supplementary Figure 6. Aggregation kinetics with Thioflavin-T fluorescence assay and different kinetic models, and Ingenuity Pathway Analysis of RNAseq from isolated microglia. A** Thioflavin-T (ThT) fluorescence intensity by time showing aggregation of 5 μM Aβ with and without increasing concentrations of Gal3 (n=8/group). Light color filling shows standard deviation. Solid lines represent fits where either k_2_ (1), k_+_ (2) or k_n_ (3) are fitted while the rate constants observed for Aβ alone are kept as global constants. **B** Thioflavin-T (ThT) fluorescence intensity by time showing aggregation of 10 μM Aβ with and without 20 μM of Gal3, run to assess the timepoints used in Fig. 5B and C (marked with red lines). Light color filling shows SD. Rationale behind selected timepoints goes as it follows: 5 min = before ThT fluorescence; 15 min = T½ of Aβ alone; 30 min = T½ of Aβ+Gal3; 80 min = after fibrillations are complete. **C** Ingenuity Pathway Analysis (IPA)’s Regulator Effects prediction of inflammatory response from isolated APP and APP-Gal3KO microglia. **D** IPA’s network display of TNF as upstream regulator from isolated APP and APP-Gal3KO microglia. In C and D, red and green indicate observed upregulation and downregulation in APP-Gal3KO microglia respectively; while orange and blue indicate predicted activation and inhibition in APP-Gal3KO microglia respectively.

**Supplementary Table 1**. **Neuropathological findings and subject information of postmortem samples.**

| **Diagnose** | **NBB no.** | **ID** | **Age** | **Gender** | **Abeta** | **NFT** | **DLB** |
| --- | --- | --- | --- | --- | --- | --- | --- |
| AD | 15-37 | AD5 | 72 | M | C | 6 | 0 |
| AD | 15-36 | AD8 | 82 | M | C | 5 | 0 |
| AD | 16-105 | AD4 | 68 | M | C | 5 | 0 |
| AD | 16-66 | AD3 | 94 | M | C | 5 | 0 |
| AD/DLB | 15-98 | AD7 | 64 | M | C | 6 | 6 |
| AD | 16-48 | AD2 | 84 | F | C | 4 | 0 |
| AD | 16-92 | AD1 | 81 | F | C | 4 | 0 |
| AD/DLB | 15-54 | AD9 | 90 | F | B | 2 | 5 |
| AD/DLB | 15-46 | AD6 | 90 | F | C | 6 | 5 |

Including diagnosis, Biobank number, internal ID, age, gender, Aβ score, tau tangles, Lewy body presence. AD = Alzheimer’s disease, DLB = Dementia with Lewy body, NBB = Netherlands brain bank, ID = Identification, M = Male, F = Female, NFT = Neurofibrillary tangles. Abeta score: indicates AB phase and ranges from A (no Aβ plaques) to C (final Aβ phases 4-5). NFT score: indicates Braak NFT stage and ranges from no tangles (0) to final Braak stage VI (6). DLB score: indicates presence of Lewy bodies throughout Braak staging (1-6).

**Supplementary Table 2**. **Full list of Gal3 interacting partners within the human proteome, ranked by pDock scores, using AlphaFold2 (FoldDock pipeline)**.

**Supplementary Table 3. Full list of gene expression results in APP-Gal3KO microglia compared to APP microglia**

**Supplementary Table 4**. **List of IFN-α and MGnD gene-sets enrichment analyzed with GSEA**.

**Supplementary Table 5. Full list of upstream regulators predicted by QIAGEN’s IPA from APP-Gal3KO microglia versus APP microglia**.

| **Primary Antibodies** | | |
| --- | --- | --- |
| Mouse anti-β-Amyloid 1-16 (6E10) | BioLegend | Cat: 803001; RRID:AB_2564653 |
| Mouse anti-b-Amyloid 40/42 (MOAB2) | Biosensis | Cat: M-1586-100; RRID:AB_2492497 |
| Goat anti-Galectin-3 | R&D Systems | Cat: AF1197; RRID:AB_2234687 |
| Sheep anti-TREM-2 | R&D Systems | Cat: AF1729; RRID:AB_354956 |
| Mouse anti-TLR-4 | Santa Cruz Biotechnology | Cat: sc-293072; RRID:AB_10611320 |
| Mouse anti-β-actin horseradish peroxidase conjugated | Sigma-Aldrich | Cat: A3854; RRID:AB_262011 |
| Mouse anti-actin | Sigma-Aldrich | Cat: A3853; RRID:AB_262137 |
| Rat anti-mouse CD45-AlexaFluor 488 | BioLegend | Cat: 103122; RRID:AB_49353 |
| Rat anti-mouse CD11b-PE/CY7 | Biosite | Cat: ASB-8VX2T6-0.1 |
| CD11b-APC | Immunostep | Cat: M11BA; RRID:AB_11140720 |
| CD45-PE | Immunostep | Cat: M45PE; RRID:AB_11139704 |
| Rabbit anti-amyloid fibrils OC | Millipore | Cat: AB2286; RRID:AB_1977024 |
| Goat anti-Cathepsin B | R&D Systems | Cat: AF965; RRID:AB_2086949 |
| Rat anti-LAMP1 | Developmental Studies Hybridoma Bank | Cat: 1D4B; RRID:AB_2134500 |
| Mouse anti-α-tubulin | Sigma-Aldrich | Cat: T9026; RRID:AB_477593 |
| Mouse anti-Amyloid-beta peptide (BAM10) | Sigma-Aldrich | Cat: A3981; RRID: RRID:AB_1078153 |
| Rabbit anti-Iba1 | Wako | Cat: 019-19741; RRID:AB_839504 |
| Rat anti-CD68 | Invitrogen | Cat: 14-0681-82; RRID:AB_2572857 |
| Rat anti-LAMP1 | DSHB | Cat: 1D4B-S; RRID: RRID:AB_2134500 |
| **Secondary antibodies** | | |
| Horse anti-mouse IgG horseradish peroxidase conjugated | Vector Laboratories | Cat: PI-2000; RRID:AB_2336177 |
| Horse anti-goat IgG horseradish peroxidase conjugated | Vector Laboratories | Cat: PI-9500; RRID:AB_2336124 |
| Rabbit anti-sheep IgG horseradish peroxidase conjugated | Invitrogen | Cat: 61-8620; RRID:AB_88263 |
| Nanogold Rabbit anti-goat IgG | Nanoprobes | Cat: 2005; RRID:AB_2617133 |
| Donkey anti-rabbit Alexa Fluor 488 | Invitrogen | Cat: A-21206; RRID:AB_2535792 |
| Donkey anti-mouse Alexa Fluor 647 | Invitrogen | Cat: A-31571; RRID:AB_162542 |
| Donkey anti-goat Alexa Fluor 647 | Invitrogen | Cat: A-21447; RRID:AB_2535864 |
| Donkey anti-rabbit Alexa Fluor 647 | Invitrogen | Cat: A-31573; RRID:AB_2536183 |
| Donkey anti-mouse Alexa Fluor 568 | Invitrogen | Cat: A10037; RRID:AB_11180865 |
| Donkey anti-rat Alexa Fluor 647 | Invitrogen | Cat: A48272; RRID:AB_2893138 |
| Donkey anti-sheep Alexa Fluor 568 | Invitrogen | Cat: A-21099; RRID:AB_2535753 |

**Supplementary Information References**

19.	P. E. Harshil Patel, Jonathan Manning, Maxime U Garcia, Alexander Peltzer, Rickard Hammarén (2024) nf-core/rnaseq: nf-core/rnaseq v3.18.0 - Lithium Lynx. (Zenodo, Zenodo).

26.	H. Wickham, *ggplot2: Elegant Graphics for Data Analysis* (Springer-Verlag New York, 2016).

27.	A. Kramer, J. Green, J. Pollard, Jr., S. Tugendreich, Causal analysis approaches in Ingenuity Pathway Analysis. *Bioinformatics* **30**, 523-530 (2014).
